# Refined Enterotyping Reveals Dysbiosis in Global Fecal Metagenomes

**DOI:** 10.1101/2024.08.13.607711

**Authors:** Marisa Isabell Keller, Suguru Nishijima, Daniel Podlesny, Chan Yeong Kim, Shahriyar Mahdi Robbani, Christian Schudoma, Anthony Fullam, Jonas Schiller, Ivica Letunic, Wasiu Akanni, Askarbek Orakov, Thomas Sebastian Schmidt, Federico Marotta, Anja Telzerow, Diënty HM Hazenbrink, Rajna Hercog, Stefanie Kandels, Robert Schierwagen, Sabine Klein, Maximilian Joseph Brol, Wenyi Gu, Frank Erhard Uschner, Maria Papp, Wim Laleman, Debbie Shawcross, Minneke J. Conraad, Rajiv Jalan, Katrine Holtz Thorhauge, Stine Johansen, Maja Thiele, Aleksander Krag, Jonel Trebicka, Michael Kuhn, Thea Van Rossum, Peer Bork, the MICROB-PREDICT consortium

## Abstract

**Background:** Enterotypes describe human fecal microbiomes grouped by similarity into clusters of microbial community composition, often associated with disease, medications, diet, and lifestyle. Numbers and determinants of enterotypes have been derived by diverse frameworks and applied to cohorts that often lack diversity or inter-cohort comparability.

**Results:** To overcome these limitations, we selected 16,772 fecal metagenomes collected from 38 countries to revisit the enterotypes using state-of-the-art fuzzy clustering and found robust clustering regardless of underlying taxonomy, consistent with previous findings. Quantifying the strength of enterotype classifications enriched the enterotype landscape, also reflecting some continuity of microbial compositions. As the classification strength was associated with the patient’s health status, we established an “Enterotype Dysbiosis Score” (EDS) as a latent covariate for various diseases.

**Conclusion:** This global study confirms the enterotypes, reveals a dysbiosis signal within the enterotype landscape, and enables robust classification of metagenomes with an online “Enterotyper” tool, allowing reproducible analysis in future studies.

**Graphical Abstract:** 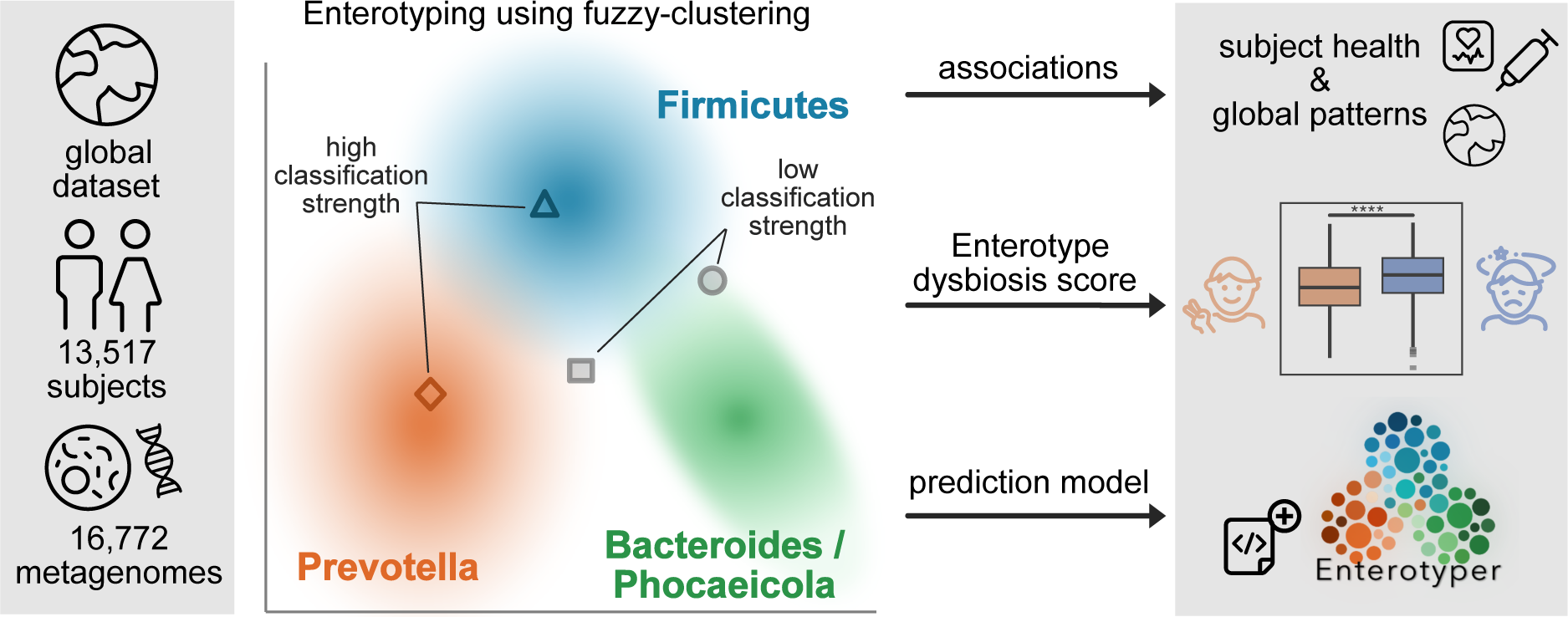

## Introduction

The human fecal microbiota has emerged as a critical axis of our health^1^. Unique in its composition for each individual^2^, the microbiota is inherently shaped by microbial interactions and external influences^3^. Environmental and host factors, including genetics, physical activity, diet, and medications, contribute to the microbiome’s variability, as do both specific and unspecific effects that appear in many diseases^4,5^. While specific alterations enable microbial signatures as biomarkers, broader microbiome dysregulations commonly called ‘dysbiosis’ tend to be more unspecific and are typically characterized by an unfavorable compositional divergence from healthy reference cohorts^6^. However, the definitions and causes of dysbiosis vary, and its implications are still poorly understood^7^.

To simplify the complex structure of the fecal microbiome, taxonomic profiles have been grouped into distinct, reproducible microbial community clusters (often at the genus level) called ‘enterotypes,’ which are typically named by their dominant taxa characterizing each cluster^8^. While most studies consistently detected enterotypes with high relative abundance of the *Prevotella* or *Bacteroides* genera, others reported communities dominated by Clostridiales^9^, Firmicutes^10,11^, or *Ruminococcaceae*^8^. Although the exact count and constitution partially differed between studies, enterotypes have been valuable in clinical research contexts for stratifying study populations based on enterotype-specific phenotypes^12,13^, responses to treatments^9,14^, and dietary interventions^15^.

The methods previously used to define enterotypes, as well as the size and diversity of underlying populations, have varied greatly, making comparisons across studies difficult. Enterotypes were first established in 2011 based on partitioning around medoid (PAM) clustering of 33 samples from three continents, from which three clusters emerged, each dominated by either *Bacteroides, Prevotella,* or *Ruminococcus*^8^. These enterotypes were supported by Dirichlet multinomial mixtures (DMM)^16^ modeling on 1106 metagenomes from Belgian individuals^17^ and a meta-cohort of 785 individuals from the USA, Europe, and China^10^.

These studies do not rule out other configurations of enterotypes, as the size of the study cohort was still below the reported lower limit for robust metagenome-wide association studies^18^. Indeed, similar work on 888 samples collected in France, Denmark, and Germany pointed to an additional *Bacteroides* enterotype associated with disease, inflammation, low fecal cell counts, and a high BMI^14^. Other analyses found between two and four enterotypes but relied on 16S rRNA gene sequencing data or were limited to homogenous study populations from single countries^19^.

More recent work by Tap et al. applied DMM clustering to 3,233 metagenomes, which was then used to predict enterotypes-like groups (called ‘branches’ by the authors) and study their trajectory using the PHATE algorithm^20^. Frioux et al. leveraged bacterial co-occurrence patterns to identify enterotype-like groups (called ‘enterosignatures’ by the authors) in 5,230 metagenomes but did not include samples from non-Westernized populations^21^. Both studies found clusters that largely overlapped with the initially proposed enterotypes.

A recurring criticism of the concept of enterotypes was the initial assumption of discrete clusters of microbiome states. It has been argued that the microbiome follows a continuous gradient of inversely related *Prevotella* and *Bacteroides* abundances^22,23^. Thus, a hybrid model has been proposed that may better reflect reality^10^, where low-density gradients connect high-density clusters (enterotypes).

Here, we developed such a quantitative model by revisiting the enterotype concept using fuzzy k-means (FKM) clustering^24^ on a curated, comprehensive, diverse, and, to date, the most extensive set of 16,772 metagenomes collected globally from 38 countries. Using a rarefaction approach, we observe the enterotype classification becoming more stable with an increasing number of samples. Leveraging FKM properties, we quantify the enterotype classification strength for each sample, effectively accounting for the inherent continuity of the microbiome composition landscape. Following the Anna Karenina principle for microbiomes^25^, which postulates that “all healthy microbiomes are similar; any dysbiotic microbiome is dysbiotic in its own way,” we observe that microbiomes with lower enterotype classification strength are associated with a variety of diseases, leading us to propose the ‘Enterotype dysbiosis score’ (EDS). Both, EDS and the enterotype classifications, should inform clinical decision making and are publicly accessible through the “Enterotyper” tool at https://enterotype.embl.de.

## Results

### Development of robust and global enterotypes

We assembled a global dataset of 16,772 metagenomes from 129 studies (Table S1) collected from 13,517 individuals in 38 countries (Figure 1A). This population had a mean BMI of 24 (SD 5.34, n=5,036), a mean age of 40 years (SD 17.15, n=7,432), ranging from 5 to 70 years, and included 32 reported diseases and medical conditions.

**Figure 1:**
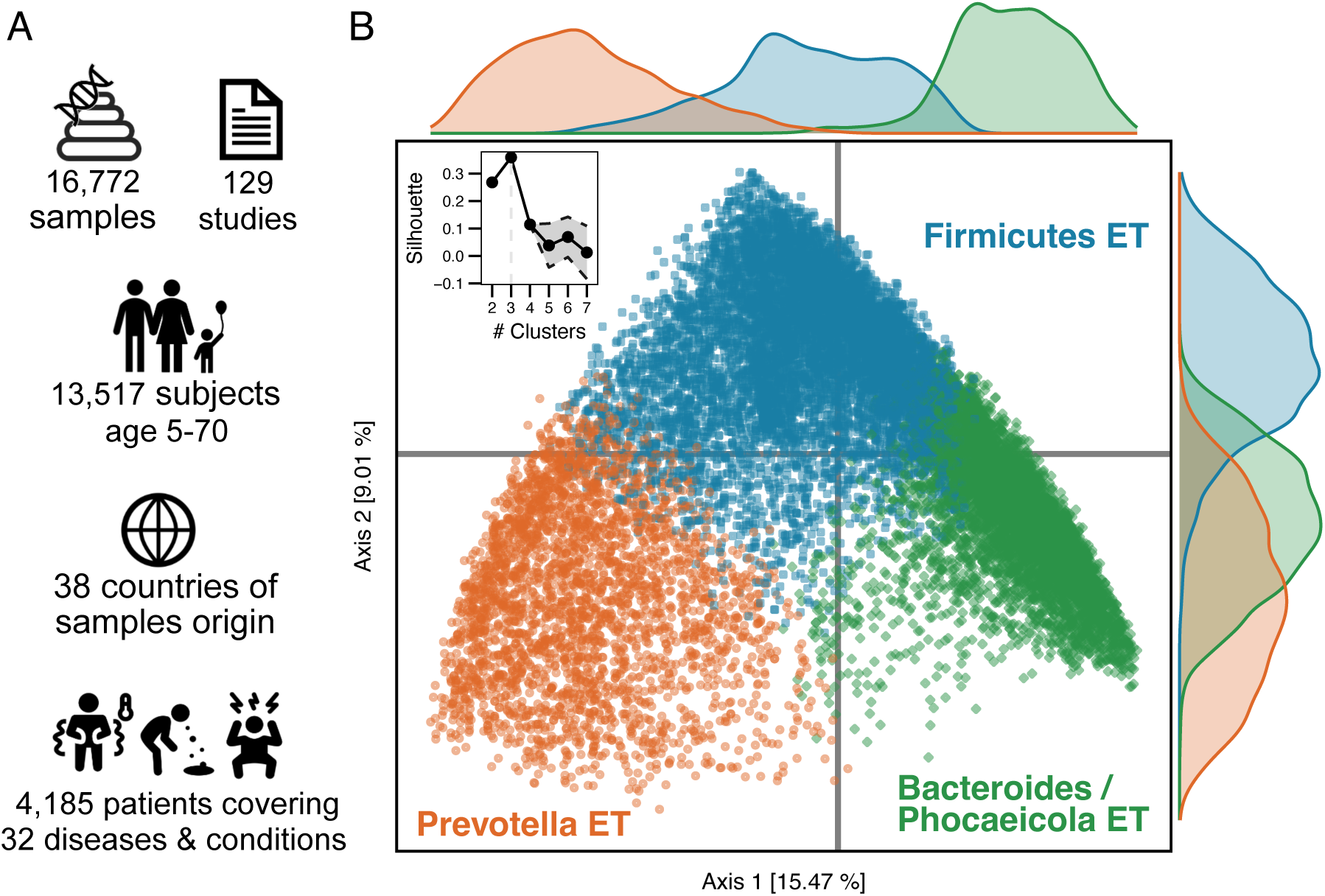
Development of robust and global enterotypes. **A.** Data overview on our global dataset. **B.** PCoA generated from the JSD matrix. Each sample is colored based on its enterotype classification. Density plots indicate the distribution of points along the first two principal components for each enterotype. The inset plot in the upper left corner displays the silhouette width and grey confidence intervals for 20 repetitions of the algorithm.

To assess the applicability of enterotyping on the global dataset and compare the results with previous work, we applied two commonly used clustering methods, PAM and DMM^10^. DMM clustering did not indicate an absence of clusters or an exact number of clusters. Instead, the optimal number of clusters was the maximum attempted, 15 or 50 (Figure S1A-B). We benchmarked the effect of dataset size and heterogeneity and found that increasing the number of samples resulted in higher numbers of clusters (Figure S1C-D). This is in line with previous DMM clustering on 3,233 samples resulting in 24 clusters^20^. Thus, DMMs alone are unlikely to reduce this dataset’s complexity into simple models as they introduce too many degrees of freedom.

Using PAM clustering, the silhouette width suggested that two, three, and four enterotypes described the dataset well, with slightly higher support for two enterotypes (Figure S2A). Further evaluation with the CH index predicted the presence of three clusters (Figure S2C). According to a PERMANOVA analysis, batch effects between studies are the predominant factor driving microbial variance across samples, explaining 10.5% of the total variance (adjusted q < 0.01; Table S2). Therefore, we wanted to estimate the effect of integrating multiple studies into a single dataset on predicting the optimal cluster number. We performed a rarefaction analysis by randomly selecting increasing subsets of studies. As the number of samples and studies increased, the optimal cluster number stabilized, with three clusters identified by the CH-index and two by the silhouette width (Figure S2B+D). These models differ because the Firmicutes and Bacteroides enterotypes are separate in the 3-cluster model and combined in the 2-cluster model (Figure S3). Lumping samples into one larger cluster loses information but splitting them into two clusters may overestimate their distinctiveness. It is, therefore, necessary to quantify enterotypes, taking the inherent continuity of microbiome composition into account. Fuzzy clustering allows overlapping clusters and quantifies the enterotype classifications, which results in more nuanced and stable enterotyping across diverse and large datasets.

### Fuzzy *k*-means (FKM) clustering confirms and enriches enterotype landscape

We applied the fuzzy *k*-means (FKM) algorithm^24^ and identified three enterotypes as the best approximation to describe the microbial compositional landscape (Figure 1B, Figure S4B). The rarefaction analysis for FKM, similar to the one for PAM clustering, demonstrated that an increased number of samples and studies stabilized the prediction (Figure S4A), emphasizing the importance of a large, global, and balanced study population for accurately calculating enterotypes given the heterogeneous nature of the microbiome. Larger cohort sizes are more likely to contain a critical mass of samples belonging to each enterotype. In contrast, analysis based on a single limited cohort may result in clusters reflecting study-specific effects rather than corresponding to globally consistent enterotypes. The wide range of different enterotype proportions in the individual studies (Figure 2A, Figure S5) supports this hypothesis that clustering on single cohorts would not result in the same enterotype classifications, as cluster methods are usually biased towards detecting equally sized clusters^26^.

**Figure 2:**
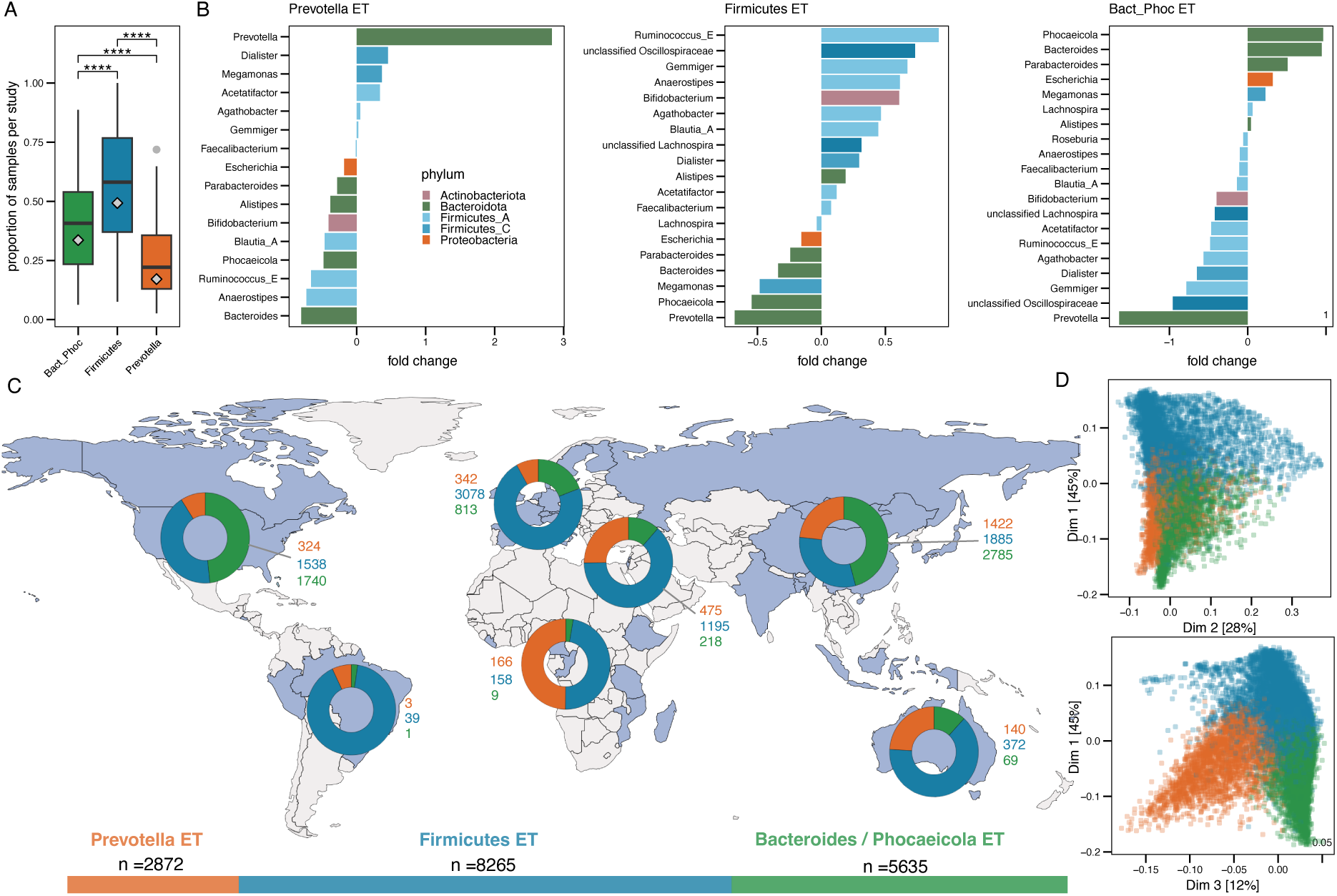
Fuzzy k-means (FKM) clustering confirms and enriches the enterotype landscape. **A.** The proportion of samples classified into enterotypes for each individual study, with the proportion in the total cohort indicated by grey diamonds. **** p ≤0.0001 **B.** Fold change from differentially abundant genera between samples classified and not classified into each enterotype. Only the top significant genera after Benjamini-Hochberg p-value correction for multiple hypotheses are shown. **C.** World map showing the global enterotype distribution with the numbers of samples classified into each enterotype reported next to the donut plots. The interactive version of that map can be explored on our webpage https://enterotype.embl.de/, which includes more information about the included studies, countries, samples, and country-wise enterotype distributions. The bar plot at the bottom shows the Enterotype classification distribution for the whole global dataset **D**. Multidimensional scaling plots with dimensions one and two (upper plot) and dimensions one and three (lower plot), calculated on the functional profiles with relative abundances of KEGG orthologs for each sample. The colors of the points correspond to the enterotypes, as reported in panel C.

One global enterotype we identified was associated with high *Bacteroides* and *Phocaeicola* abundance (‘Bacteroides/Phocaeicola enterotype’, Figure 2B). The latter genus was recently split off the *Bacteroides*^27^. Their relative abundances correlate (R=0.64, p < 2.2e-16, Pearson correlation, log10-transformed data) and jointly drive this enterotype. *Prevotella* dominated a second enterotype, and the third enterotype was driven by several genera, mainly belonging to the Firmicutes phylum (e.g., *Ruminococcus_E, Gemminger, Anaerostipes, Dialister, Blautia*) and was therefore named the ‘Firmicutes’ enterotype. These taxonomic annotations are all based on the GTDB^27^ taxonomy, which is increasingly used in metagenomic studies instead of the traditional NCBI^28^ classification. We compared both taxonomies to assess their influence on enterotypes. The silhouette width showed consistent results (Figure S4E), and enterotype classification resulted in 94.8 % (n=15,852) of the samples being classified into the same enterotype regardless of the underlying taxonomy.

FKM calculated the strength with which the microbiomes belong to each enterotype by reporting the consistency of enterotype classifications over multiple cluster iterations. This “classification strength” offers new perspectives on the microbiome landscape, presenting a hybrid design between distinct enterotype clustering and the gradient concept that considers the continuous nature of fecal microbial composition. We found that FKM classifies 87.5% (n=14,675) of samples into the same three enterotypes as assigned by PAM clustering (Figure S4C) while reporting lower classification strength for the samples that have different classifications across the two models (Figure S4D).

### Enterotypes built on taxonomic profiles reflect microbial metabolic differences

We aimed to quantify the extent to which enterotypes, as preferred taxonomic compositional states, vary in their functional profiles and, consequently, their impact on the host. Using a dimensional reduction method (MDS), the functional profiles for each sample showed a remarkable alignment with enterotype classifications (Figure 2D). The Firmicutes enterotype separates from the others across the first dimension, while the Bacteroides/Phocaeicola and Prevotella enterotype separates across the third dimension. We compared the differential abundance of KEGG-Orthologs between enterotypes (Figure S6) and found, for example, pronounced differences in carbohydrate uptake mechanisms. The Bacteroides/Phocaeicola and Prevotella enterotypes are characterized by specialized outer membrane proteins for starch uptake^29^. In contrast, the Firmicutes enterotype shows an ATP-binding cassette (ABC system) enrichment associated with glycan uptake^30^.

### Robustness of enterotypes across methodology in integrated cohorts

Recent studies of combined cohorts used different approaches to summarize the complexity of the human microbiome into higher-level community groups^20,21^, which we wanted to compare to the FKM-derived enterotypes. Tap and colleagues used DMM on 3,233 metagenomes, subsequently imposing the clustering on around 15,000 additional samples followed by hierarchical clustering^20^, while Frioux and colleagues focused on bacterial co-occurrence patterns^21^. The high overlap of samples between Tap et al. and this study enabled us to link the sample classifications directly. To include data from Frioux et al. on the intersection, we used their online prediction tool on our cohort. The results across the different approaches are consistent, identifying three major clusters associated with *Bacteroides, Prevotella,* or Firmicutes species (Figure 3A & S7A). Between Tap et al. and this study, 4,132 samples (75,5 %) are consistent in their enterotype classification, while Frioux et al. and this study agree in 11,551 (72,3 %) samples. Samples showing inconsistent classification were associated with a lower FKM-derived classification strength (Figure 3B & S7B), reinforcing the substantial overlap between the sample classification for central enterotype samples.

**Figure 3:**
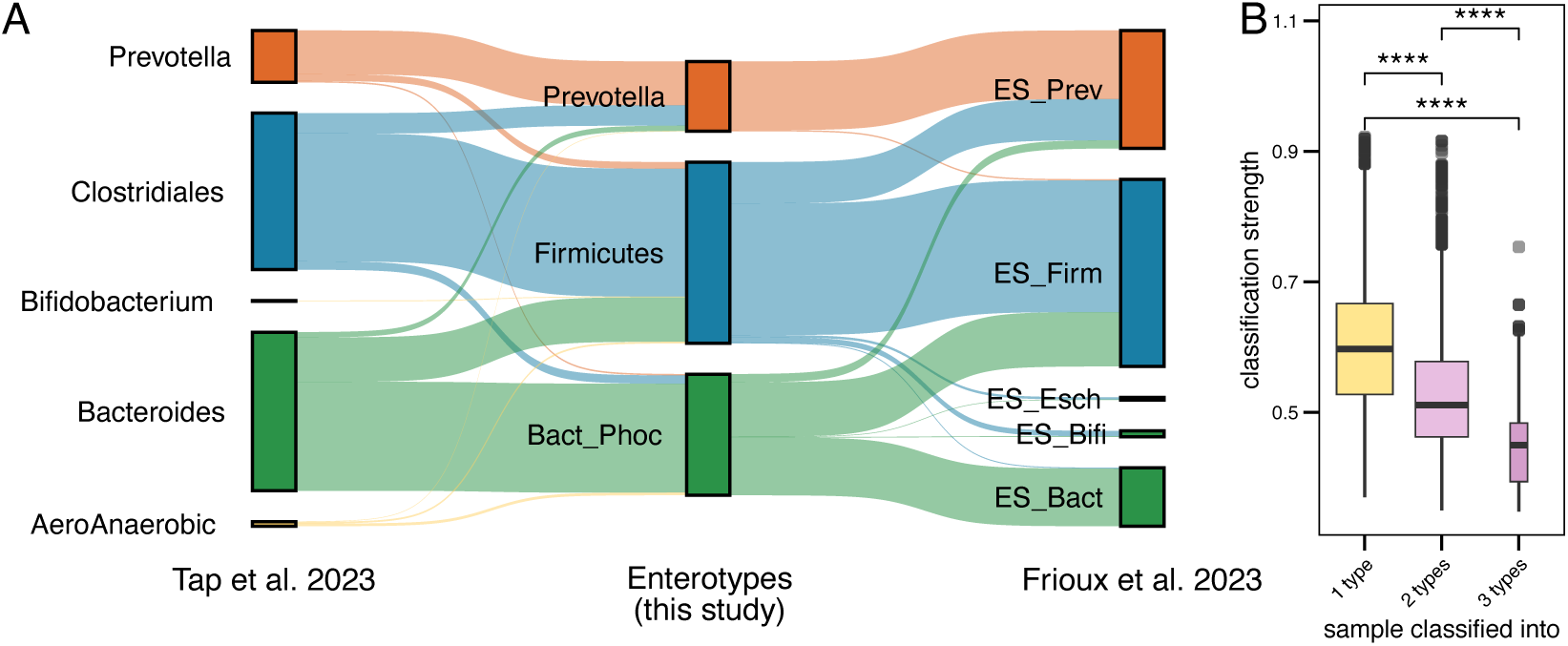
Robustness of enterotypes across methodology in integrated cohorts. **A.** Comparison of the enterotype classifications for the overlapping samples between this study and Tap et al. 2023 and Frioux et al. 2023 (n = 5,668), which used alternative approaches to define compositional groups in the human fecal microbiome. **B.** The stability in the enterotype classification among the different studies in panel A is associated with FKM-derived classification strength, showing lower strength for samples with changing classifications. 2,749 samples are stable, 2,430 samples change once, and 295 samples change twice in their enterotype classification.

Additional groups in Tap et al. and Frioux et al. show only a small overlap with our dataset as they are mainly found among young children, which motivated us to contextualize the mature human enterotypes from the FKM on our global dataset. We combined our dataset with other human gut microbiome samples from infants (< 5 years) and older people (>70 years), as well as samples from different human body sites and other animal host-associated habitats (total n = 49,762). Using a dimensional reduction method (UMAP)^31^, we observed a clear distinction between the human gut microbiomes of adults and infants and those of other human and non-human habitats (Figure S8). While different dimensionality reduction methods were used to observe the host and body site separation,^32,33^ we also found a reflection of the enterotype classifications. This highlights the distinct enterotype signal in the human gut microbiome even when clustered with vast heterogeneous samples.

### Stability of enterotypes and associations with demographic measures and diseases

Our dataset is a rich resource for testing the temporal stability of enterotype classification, as it contains 651 subjects with multiple samples within one year (3,853 samples, Figure S9A). We used a Markov chain model to assess the stability of classifications (Figure 4A; examples in Figure S9B-C) and found that 57% (n=362) of the subjects showed high stability in their enterotypes. Among all enterotype changes, the most frequent is to the Firmicutes enterotype, starting from the Bacteroides/Phocaeicola (186 changes, 28%) and Prevotella (116 changes, 17%) enterotypes. Transitions between the Prevotella and Bacteroides/Phocaeicola enterotypes were rare (49 changes in both directions, 7%) and possibly induced by microbiome perturbations. Many subjects with changes in enterotypes suffer from inflammatory bowel disease (n=55), a condition that is known to increase microbial variability^6^, or took part in a probiotic intervention study (n=32)^34^.

**Figure 4:**
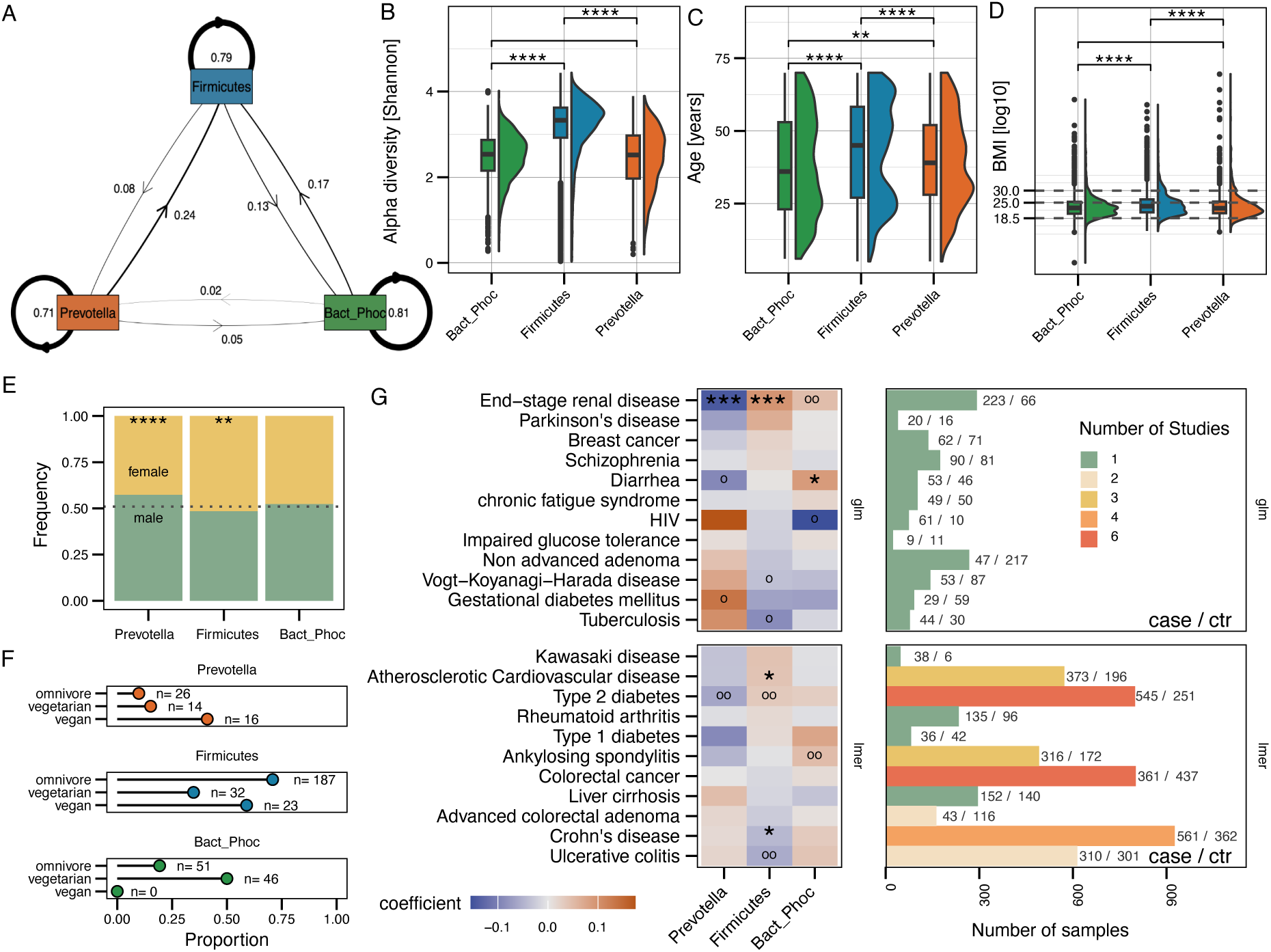
Stability of enterotype and associations with demographic measures and diseases. **A.** A Markov chain model estimation of the likelihood of transitioning between enterotypes for subjects who provided two or more samples within one year. **B. – D.** Shannon diversity, age, and BMI comparisons between the samples classified into different enterotypes. The cohort was filtered for all samples with information on age (n=7,432) and BMI (n=5,036). Differences between the groups were calculated using the Wilcoxon test. *p ≤ 0.05, **p ≤ 0.01, ***p ≤ 0.001, ****p ≤ 0.0001 **E.** Distribution of 7,973 subjects with sex information in the metadata. Using a chi-squared goodness-of-fit test, we checked if the observed ratio for each enterotype deviated from the expected ratio in our cohort (0.49:0.51, dotted line). **p ≤ 0.01, ****p ≤ 0.0001. **F.** The proportion of samples from subjects reporting a dietary pattern, such as omnivore, vegetarian, and vegan, classified into the enterotypes. The numbers of samples are noted in the plot. **G.** Associations between diseases or symptoms and the probability for the corresponding enterotype were tested using generalized linear models (glm) with additional adjusting for random effects using linear mixed effect models (lmer) to account for longitudinal sampling and study effects where appropriate. A positive coefficient (orange) refers to a positive correlation between enterotype affiliation and disease or condition for a given sample. In contrast, a negative coefficient (blue) means that samples from subjects with this disease or condition are less likely to be classified to this enterotype. For each disease or condition, the models were calculated on available disease and control studies, and the number of cases and controls and the number of studies are reported in the bar plot. ° p ≤ 0.05, ^oo^ p ≤ 0.01, * q ≤ 0.05, *** q ≤ 0.001; q=adjusted p-values using Benjamini-Hochberg correction.

After having established a robust global enterotype framework that is independent of underlying taxonomy and study effects as well as stable over time in an individual, we then adequately explored the relationships between enterotypes and host phenotypes. We found the Firmicutes enterotype microbiomes had higher species-level alpha diversity, and the subjects were often older, female, and had higher BMIs (Figure 4B-E). Male subjects were overrepresented in the Prevotella enterotype (Figure 4E). Our dataset includes 395 samples from five studies with documented dietary patterns, which revealed a significant association between diet (omnivore, vegetarian, vegan) and enterotypes (chi-squared test, p = 2.3e-15; Figure 4F). Plant-based diets were most associated with the Prevotella enterotype. Only 10% of omnivores (n=26) were classified as the Prevotella enterotype, compared to 15% (n=14) of vegetarians and 41% (n=16) of vegans. Notably, no vegan had microbiomes classified to the Bacteroides/Phocaeicola enterotype. This adds to previous hypotheses based on smaller datasets implying that dietary habits at least contribute to enterotype stratification^35^.

To explore the relationship between disease and symptoms with the enterotypes, we used linear mixed-effect models (lmer), adjusting for multiple studies and samples, and generalized linear models (glm) for single-study, non-repeated sampling data (Figure 4G). Instead of limiting the analysis to rigid enterotype classifications, we used the classification strength of each sample for each enterotype to provide a more nuanced analysis. We only analyzed the study’s internal samples to use matched cases and controls. End-stage renal disease was associated positively with the Firmicutes enterotype (glm, coefficient = 0.096, q = 3.2e-05) and negatively with the Prevotella enterotype (glm, coefficient = −0.14, p = 8.8e-03), which is consistent with previous findings^36^. Diarrhea was associated with the Bacteroides/Phocaeicola enterotype (glm, coefficient = 0.088, q = 2.2e-02), and cardiovascular disease showed associations with the Firmicutes enterotype (lmer, coefficient = 0.04, q = 1.9e-02).

### Enterotype dysbiosis scores (EDS) are linked to subjects’ disease status

Besides the associations of specific diseases or symptoms with the enterotypes, we wondered to what extent samples from diseased individuals generally fit into the enterotype landscape. FKM clustering quantifies how well a microbiome fits in each enterotype using the consistency of classification across cluster iterations in the process of defining the best cluster number. This is reflected visually in a PCoA, where samples central to an enterotype have more robust classification than those between enterotypes (Figure 5A). We hypothesized that samples with lower classification strength are more dysbiotic and developed a classification strength-based Enterotype dysbiosis score (EDS). A higher EDS indicates a higher deviation from enterotype centers, which translates to dysbiotic microbiomes being located somewhere in the space “between” or “beyond” enterotypes.

**Figure 5:**
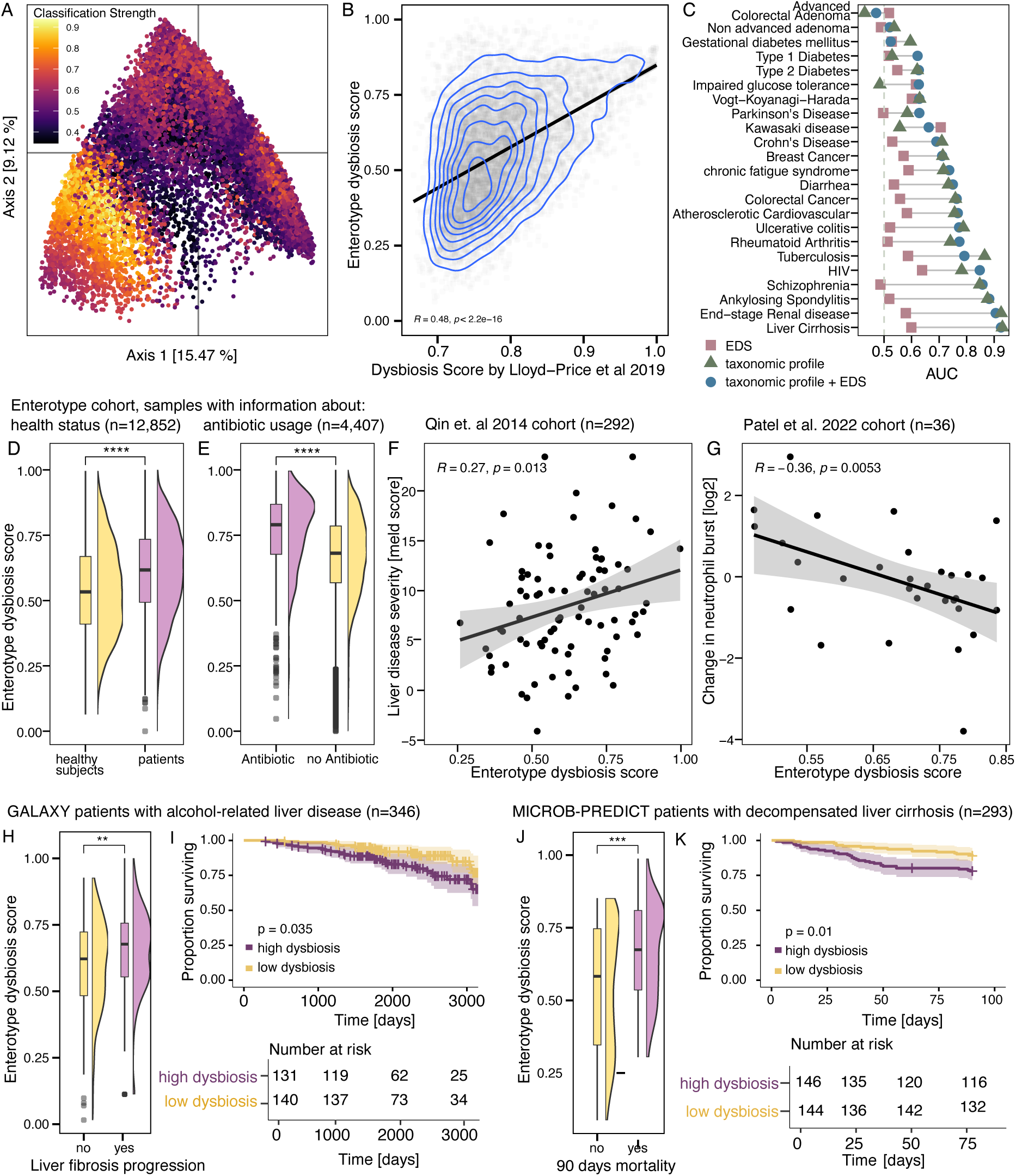
The enterotype dysbiosis scores (EDS) are linked to subjects’ disease status. **A.** Principal coordinates analysis (PCoA) plot based on JSD from genus-level GTDB taxonomic profiles. Each sample is colored based on its enterotype classification strength. The samples with higher probabilities represent the “centers” of the enterotypes in the two-dimensional space. **B.** AUCs from disease classification models based on taxonomic profiles, EDS alone, and models with combined EDS and taxonomic features. The models were trained on study intrinsic case and controls. **C.** Samples from subjects with a reported disease (n = 3,399) showed a higher EDS than those from subjects with no reported disease (n = 8,672). **D.** Subjects who reported antibiotic use (n=1,038) showed higher EDS. **E.** Spearman correlation of the EDS with the MELD (model of end-stage liver disease) score, reported for 181 subjects in Qin et al.^65^, as an estimate of liver disease severity. **F**. Alcohol-related liver disease (ALD) patients showing higher rifaximin-⍺ treatment response (Change in neutrophil burst) had a higher baseline EDS score. **G**. ALD patients who progress in disease within 8 years follow-up show higher dysbiosis at baseline than patients who stayed stable or improved. **H.** Kaplan-Meier plot showing that EDS can be used to stratify ALD patients for their probability of surviving. The p-value was calculated using a log-rank test. **I.** Patients with decompensated cirrhosis who died within the 90-days follow up show higher EDS than surviving patients. **J.** EDS can stratify surviving and non-surviving patients with decompensated cirrhosis. Samples from panel G-K were not included in the global enterotype dataset but the enterotype and EDS have been predicted using the highly accurate prediction models (Figure 6). ** p ≤ 0.01, *** p ≤ 0.001, **** p ≤ 0.0001.

EDS showed a significant correlation with a previously established dysbiosis score, which defines dysbiotic samples as different to a healthy reference set^6^ ( Figure S10A). Alpha diversity as a common measure of gut health^37^ showed a negative relationship with EDS (Figure S10B). Healthy subjects had lower EDS (Figure 5C, Figure S10C). Twenty out of 32 medical conditions, including colorectal cancer, Crohn’s disease, and ulcerative colitis, were associated with higher EDS (Figure S11). Most of these diseases have been previously linked to alterations in microbiome composition^5,38,39^. The most dramatic shift in EDS was seen in patients with a reported *Clostridium difficile* infection. This could be explained by repeated antibiotic usage of these patients, which is associated with a higher EDS (Figure 5D).

We used the same set of case-control studies as above (Figure 4G) to assess the relative contributions of dysbiosis and specific microbiome changes on disease classification. For each disease, we evaluated how well a model can distinguish patients from controls from EDS alone, the taxonomic profiles, and a combination of both. As expected, taxonomic profile-based classification achieved higher AUCs for most of the diseases (Figure 5B), indicating the presence of disease-specific signatures. However, EDS outperformed taxonomic profile for advanced colorectal adenoma, impaired glucose tolerance, and Kawasaki disease, indicating that these diseases cause more unspecific perturbations to the gut microbiome. Adding EDS to the taxonomic features improved the prediction for tuberculosis, Parkinson’s disease and gestational diabetes.

To demonstrate the clinical relevance of the EDS, we analyzed cohorts with available detailed clinical metadata. In patients with liver cirrhosis^40^, EDS correlated with disease severity (Figure 5G). In another cohort, Patel et al. showed that rifaximin-α reduces inflammation via modulation of the gut microbiome^41^. We observed a significant negative correlation between EDS and the change in neutrophil oxidative burst after 30 days into the trial, the primary endpoint of the clinical trial and a marker of reduced systemic inflammation (Figure 5F). ANOVA confirmed that dysbiosis significantly explained the neutrophil burst response (F=7.23, p=0.012), independent of treatment group (F=0.003, p=0.96).

In a cohort of patients with alcohol-related liver disease (ALD)^42^ with longitudinal outcome information (mean follow up time: 5.57 ± 2.08 years), patients who progressed or died during the study had higher baseline EDS than those who remained stable or improved (Figure 5G). Similarly, in decompensated cirrhosis, baseline EDS was elevated in patients who died during follow-up (Figure 5I). In both cohorts, Kaplan-Meier analysis showed that EDS stratified survival (Figure 5H & J). The signal overlapped with established measures of liver disease severity, transient elastography (TE) and the model for end stage liver disease (MELD), respectively. However, EDS added predictive value, particularly in the low TE and high MELD groups (Figure S12).

In summary, the EDS provides a measure of dysbiosis, which could be used for further patient stratification beyond taxonomic profiling. It is independent of a particular disease or the availability of a healthy reference sample set, and accounts for the underlying microbial compositional states across various disease phenotypes.

### The online “Enterotyper” tool predicts enterotype and EDS

To facilitate enterotype classification allowing an abstract view of the state of an individual microbiome, we have developed an online prediction tool. It classifies enterotypes taking into account the strength of the clustering as well as an associated dysbiosis estimation for newly generated human fecal microbiome samples. We trained XGBoost regression models^43^ and validated them on an independent dataset (n=347, details in methods). The XGBoost models showed areas under the curve (AUC) ranging from 0.985 – 1 (Figure 6A & Figure S13) and predicted strength for the enterotype classifications correlated with the validation dataset (Figure 6B). Using the same dataset, we also validated the EDS, finding that the EDS was increased for patients with Crohn’s disease (n=71), colorectal cancer (n=46), and ulcerative colitis (n=53) compared to healthy subjects (n=177), with an AUC of 0.73 (Figure 6C-D).

**Figure 6:**
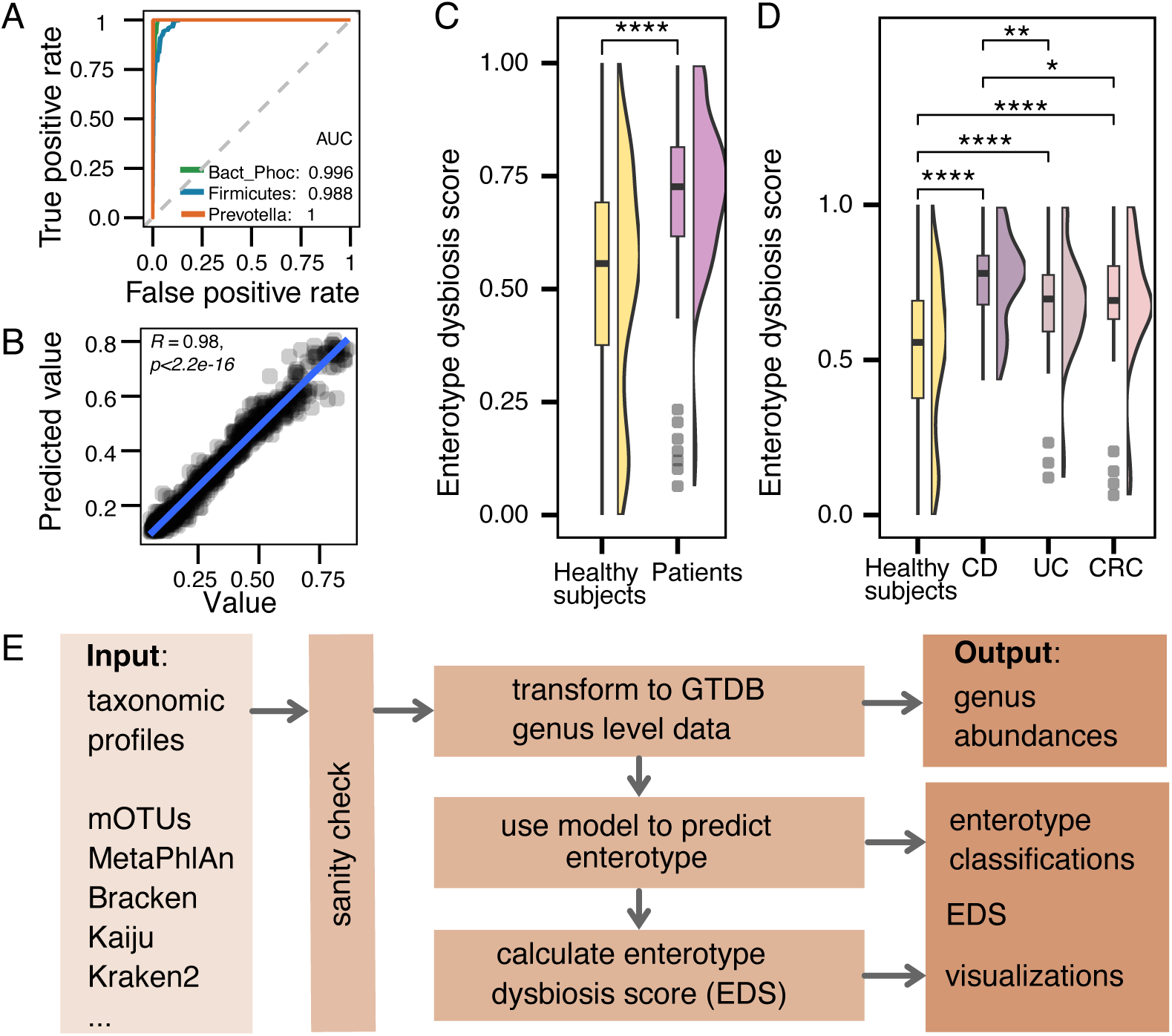
The online “Enterotyper” tool predicts enterotype and EDS. **A.** The receiver operating characteristic (ROC) curve was estimated on the validation dataset for each enterotype using the default prediction model based on the three FKM enterotypes. **B.** Correlation of predicted value and expected value for the enterotype strength for each sample in the default FKM three enterotypes estimated on the validation dataset. **C-D.** The patients (n=170) showed higher EDS than the healthy subjects (n=177) in the validation cohort. CD = Crohn’s disease, UC= ulcerative colitis, CRC = colorectal cancer. * p ≤ 0.5, ** p ≤ 0.01, **** p ≤ 0.0001. **E.** Schematic and automated workflow for the “Enterotyper” from a user perspective.

Given the overlap between the enterotype classifications between the different approaches described above and the global, comprehensive training dataset used, we have made our XGBoost models publicly available via an easy-to-use online “Enterotyper” (https://enterotype.embl.de/) to facility easy classification of a given taxonomic profile into enterotypes and estimation of the EDS (Figure 6E).

## Discussion

Enterotypes describe states of the human fecal microbiome composition associated with various factors including diseases and host lifestyle^3^, providing a useful abstraction of the complex microbiome into an easily communicated latent variable. More than a decade after the first enterotype publication^8^, we revisited the concept, quantified nuance within the model, and studied its relationship to host health and microbiome dysbiosis. A new enterotyping methodology was used on a curated set of 16,772 metagenomes, 500 times the size of the original study population. We observe that the three-enterotype landscape of Bacteroides/Phocaeicola, Prevotella, and Firmicutes-dominated communities remains the best approximation of the complex landscape of community composition.

The observation of enterotypes suggests that, across the human population, there are a few distinct alternative states into which microbial communities tend to converge. Given the symbiotic relationship between human hosts and their microbial communities, these states should promote the health of their host, at least in the sense of high-density areas. This supports a normative model for eubiosis and dysbiosis: what is typically observed in healthy individuals is the balanced, eubotic state, while rarer states and those seen in unhealthy individuals are the unbalanced or ‘dysbiotic’ states. This concept is reflected in the Anna Karenina Principle for microbiomes^25^, which argues that dysbiotic microbiomes are expected to vary widely, while eubiotic microbiomes should be more constrained. We see this reflected in our results, where healthy individuals are more likely to have a solid membership for one enterotype, and unhealthy individuals are more likely to be intermediary or completely distinct from the enterotypes. A novel aspect of the enterotype model presented in this work is that this membership strength can be quantified and translated into the Enterotype dysbiosis score (EDS). The EDS builds on a previous approach^6^ that is conceptually similar but does not consider compositional states within the eubiotic reference set. As we found associations of EDS with disease states and severity, it can be used to stratify individuals within statistical analyses and to study links between the microbiome, enterotypes, and human health.

In the enterotype landscape presented here, a second Bacteroides enterotype, previously identified through DMM in cohorts up to approximately 2,500 samples^14,44,45^, is not represented. This cluster was not seen as a distinct enterotype in our study because its samples were clustered together with the samples in the Bacteroides/Phocaeicola enterotype. This is likely due to the combined effect of DMM tending to call more clusters and the larger sample size ‘filling in the space’ between the two Bacteroides clusters. The second DMM-derived Bacteroides enterotype was previously linked to high BMI^14^, obesity^46^, metabolic disorders^13^, IBD^44^, and cardiovascular disease^45^. We acknowledge the importance of these associations and the usefulness of DMM in describing these study populations. However, the effect of reporting higher cluster numbers with higher sample numbers found in this study and by Tap et al.^20^ demonstrated the limitations of using DMM with large heterogeneous cohorts to describe broad, overarching patterns. This is an excellent example of how clustering within smaller, narrowly defined datasets might be more influenced by study- or disease-specific signals, as also shown in our rarefaction analysis, but may provide additional insights. The two goals should be clearly distinguished: one is to describe the structure of microbiomes as broadly and globally as possible (enterotypes) and the other is to define population or phenotype-specific microbiome clusters, which can be useful for investigating disease-specific and local effects. For the latter, we support the exploration of study-specific clusters by de novo clustering. For the former, we recommend that the model developed and presented here be used. To enable this, we provide researchers with the ‘Enterotyper’ tool. This provides reliable and reproducible enterotype classification of new metagenomic samples against the background of the global dataset, ensuring reproducibility and comparability between studies.

A major co-variate with enterotype is diet. The association of the Prevotella enterotype with a high-fiber diet has been previously reported^10,47^, while a diet richer in animal fat and simple sugars has been linked to the Bacteroides enterotype^10,48^. Diet is, in turn, associated with geography, and indeed the global distribution of enterotypes seen in this study overlaps with the typical dietary patterns prevalent in the different regions. North America has both the largest proportion of individuals in the Bacteroides/Phocaeicola enterotype (Figure2C) and the highest rate of animal protein consumption (83g per day per capita^49^), while Africa and Asia have the largest proportion of individuals in the Prevotella enterotype (Figure 2C) and the highest rate of more plant-based protein consumption (50g and 58g, respectively^49^). Congruently, we observed that the proportion of samples classified into the Prevotella enterotype increased with the strictness of plant-based diet (i.e., omnivore to vegetarian to vegan). In addition, the biggest change in the functional analysis of KEGG orthologs across enterotypes was linked to microbial metabolism of dietary carbohydrate compounds (starch and glycan uptake). Although diet is a major influence on the gut microbiome^17^ and enterotype classification, dietary metadata was only reported for 2% of samples (n=395) in our study population. Documenting reliable dietary data is challenging, requiring restrictive measures to record or control all food intake or relying on subjective questionnaires. To better understand effects of diet on host health and the microbiome, longitudinal studies focusing on microbial metabolism in food digestion^50^ and combined metagenomic and metabolomic studies with rich contextual data are needed^51^. Other sample biases include urban and rural differences, access to healthcare system and local dietary preferences is. While we included as much data from non-Western populations as was available, there is still a general under-representation of samples from non-Western communities in the microbiome field^52^ and we encourage more sampling efforts in non-Westernized communities.

This study advances our understanding of the diversity and constraints of the human fecal microbiome by refining the enterotype model. We increased robustness by including a much larger and more representative population and a modern taxonomy. The methods of fuzzy clustering adopted to enterotyping also allowed to utilize the correlation of classification strength and dysbiotic to develop and provide a dysbiosis estimation. Together with the easy-to-use and publicly available “Enterotyper” classification tool that allows a standardized comparison of newly generated with historic datasets.

## Methods

### Data Overview

For the global analysis, we gathered 27,674 publicly available human fecal metagenomic samples from 173 studies from the SPIRE database^53^. For our enterotype cohort, we only kept samples from subjects between 5 and 70 years old and removed samples described in original publications as deriving from “elderly” subjects. For 21 studies that provided fecal metagenomic samples from infants, we removed the samples if no age was reported for the subject. We excluded samples from fecal microbiome transplantation (FMT) studies and individuals taking antibiotics, as these interventions are likely to alter the microbiome’s composition substantially^54,55^. Samples with no reported disease status and no antibiotic treatment record (N=4,122) were not removed. If we could not link sample-specific clinical information but the study contained a general description of a disease or lack of disease for all samples in the study, we manually added that disease or lack of disease to the metadata. After filtering out samples with irregular taxonomic composition (see taxonomic and functional profiling), the remaining sample collection comprised 16,772 samples from 129 studies (see Supplementary Table 1).

The MICROB-PREDICT cohort consists of the MUCOSA-PREDICT and STOOL-PREDICT cohort^56^. Samples were gathered through a prospective, observational, multicenter study involving 48 hospitals across Europe. The institutional review board at each participating center approved the study protocol. Patients were included when a previous liver biopsy or a combination of clinical signs and findings from laboratory tests, endoscopy and ultrasonography diagnosed cirrhosis. For DNA extraction the Qiagen AllPrep Power Fecal DNA/RNA Kit (Qiagen, Hilden, Germany) was used. The generation of metagenomic sequencing libraries was done using the NEBNext Ultra II DNA library prep kit (New England Biolabs, MA, USA) and for better handling a liquid automated system (Beckman Coulter i7 Series). We aimed for a library size of 350-400 bp, along with dual index multiplex oligos. Sequencing 2x 150 bp paired-end reads was performed using an Illumina HiSeq 400 platform (Illumina, San Diego, CA, USA).

### Metagenomic data processing

#### Taxonomic and Functional profiling

Metagenomes were taxonomically profiled with mOTUs v2.5 with default parameters. Analyses were performed using two taxonomic classifications of individual mOTUs: the default taxonomy (based on the NCBI taxonomy) and a newly generated GTDB-based taxonomy (based on GTDB r202). The GTDB taxonomy implemented the reclassification of many former Bacteroides species to the genus Phocaeicola. The GTDB taxonomy was obtained by classifying genomes in proGenomes2 (corresponding to “ref_mOTUs”) using GTDB-tk and directly mapping “meta-mOTU” marker genes against GTDB marker genes, followed by rule-based and manual resolution of conflicting classifications within individual mOTUs. Six hundred and forty-six samples with more than 10 % unclassified species (“-1” fraction from mOTUs) and samples with a richness of less than ten different mOTUs were removed. The removed 646 samples are well distributed across the countries from which samples have been collected and thus not results in any significant bias against understudied populations. While the 646 removed samples represent 3.7 % of the samples, they only represent 2.5 % of the reads. The lower sequencing depth of the removed samples could hint to technical issues in the sequencing and therefore could explain the detected lower diversity and high number of unclassified reads. For a genus-level taxonomic profile, we summed the relative abundance for all mOTUs with the same genus.

Open reading frames (ORFs) were predicted using Prodigal (v2.6.3) and annotated with eggNOG-mapper (v2.1.3) against eggNOG database 5.0. Each predicted gene was multiplied by the average sequencing coverage of the contig it was found on. If a gene was associated with multiple KEGG orthologs, the average sequencing coverage count was distributed evenly between associated KEGG orthologs. A sample-wise compositional transformation yielded relative abundance values per KEGG orthologs.

#### Clustering and enterotype classification

All cluster methods used an abundance matrix on the genus level. The Enterotype clusters based on the partitioning around medoid (PAM) clustering on the samples’ Jensen-Shannon divergence (JSD) have been identified as described before^8^. In the PAM clustering, the Calinski-Harabasz (CH) index (ratio of the sum of inter-clusters dispersion and intra-cluster dispersion) together with the silhouette width (measure how similar a sample is to its cluster compared to other clusters) has been used to assess the number of clusters and cluster robustness^57^.

DMM clustering was run as reported before^16^ using the DMM R package^58^ on square root normalized relative abundances. The optimal number of clusters was evaluated by detecting a local minimum using the Laplace approximation, Bayesian information criterion (BIC), and Akaike’s information criterion (AIC). To assess the impact of sample size on DMM clustering, we randomly subsetted the global dataset into fractions of 1/2 (n=8386), 1/4 (n=4193), 1/8 (n=2096), 1/16 (n=1048), 1/32 (n=524) of the original sample size. This sub-setting was repeated three times for each fraction. To benchmark the effect of heterogeneity on the predicted optimal cluster number, we performed DMM clustering on the eight most extensive datasets within our global dataset from an increasing number of studies (number of studies mixed = 1 to 8), keeping sample size approximately constant in the mixed cohorts (number of samples = 996 to 1000). The optimal number of clusters was determined in both benchmarks using the Laplace approximation.

The fuzzy *k-*means (FKM) clustering was obtained using the *FKM* function from the fclust R package^59^ on the relative abundance matrix, and silhouette width was used as the cluster validity index to select the optimal number of clusters. The silhouette width as the clustering statistic is most comparable to the cluster statistics in the PAM clustering. The CH-index is unavailable as the within and the between the sum of squares cannot be computed in fuzzy clustering as cluster medoids are not defined. The enterotype classification for each sample was obtained from the cluster with the highest enterotype classification strength for this sample.

The PCoA was calculated using the *dudi.pco* function from the ade4 R package^60^ on the JSD matrix and the PCA based on functional profiles was calculated using *prcomp* function from the stats R package^61^.

We performed a rarefaction analysis on our cohort to assess the robustness of our PAM and FKM clustering results. This approach allowed us to investigate the impact of varying sample sizes on the stability of the predicted number of clusters. For this analysis, we selected samples from a subset of randomly chosen studies representing 30%, 40%, 50%, 60%, 70%, 80%, and 90% of the studies in our cohort. We repeated this selection process 20 times for iterations of each percentage) and performed FKM clustering for each subset in the same way as described above, utilizing silhouette width as a criterion to determine the optimal number of clusters. Additionally, we conducted PAM clustering using the CH-index and the silhouette width to evaluate the clustering quality and the best number of clusters for each subset.

For the UMAP, we compiled taxonomic profiles of animal-host-associated metagenomic samples from the SPIRE dataset ^53^. Samples with over 10% unidentified profiles were excluded from the analysis. We eliminated unidentified features for the remaining profiles and re-normalized each to ensure the total equaled one. For the remaining 49,762 samples, we calculated Bray-Curtis dissimilarity based on their GTDB-based profiles. Finally, we conducted UMAP^31^ dimensionality reduction, setting the ‘n_neighbors’ parameter to 50 and the ‘min_dist’ parameter to 0.01.

### Statistical Analysis

To demonstrate the differential abundance of genera and KEGG orthologs between enterotype clusters, we applied the Wilcoxon rank-sum test and Benjamini-Hochberg correction^62^ to adjust the p-value for multiple testing. Recurrent samples from the same subject were removed. To assess the impact of various factors on the distance matrix of the microbiome data, we performed a permutational multivariate analysis of variance (PERMANOVA) using the *adonis2* function from the vegan package in R^63^. The categorical variables of interest, such as age, sex assigned at birth, diet, or disease status, were individually tested as predictors of the microbial community structure represented in a JSD matrix.

#### Longitudinal Analysis

A Markov chain model was used to analyze the likelihood of transitions between enterotypes, providing insights into the stability and dynamics of enterotype classification and acquired using the *markovchain* R package^64^. To ensure a consistent and relevant timeframe for our analysis, we only considered samples collected within one year for each subject. In addition, we excluded the subjects with 89 samples to prevent undue influence from this individual with numerous repeated samples from the model.

#### Enterotype Associations

The associations of enterotype classifications with age, BMI, and alpha diversity (Shannon index) were calculated using the Wilcoxon rank sum test. To test differences in the gender and diet distribution, we performed a chi-squared goodness-of-fit test to determine if the observed frequency in each group deviated significantly from the ratio in our cohort.

In available case and control studies, we calculated generalized linear models (glm) using the *glm* function from the R “stats” package^61^, to model the association with reported diseases and conditions with each enterotype. For each enterotype, separate models were calculated using the strength value for this enterotype derived from the FKM clustering as a measure of enterotype affiliation. We used only the internal controls of the study to avoid artificially inducing significance through a considerable sample size of controls and to benefit from matched cases and controls that are usually selected in the case and control studies. We adjusted for random effects using the *lmer* function from the lme4 R package^65^ to account for longitudinal sampling and study effects where appropriate. Multiple testing corrections were conducted on all p-values using Benjamini-Hochberg corrections.

For the machine learning models using taxonomic profiles with and without the EDS, the SIAMCAT v2.6 machine learning toolbox was used^66^. We predicted the disease status using L1-regularized (LASSO) logistic regression model on log transformed z-score normalized data. The models were built using a 5×5 cross-validation methodology, while samples from sample subjects were kept together in either train or test set. The predictions on the test sets were used to calculate the Area under the Curve (AUC) of the Receiver Operating Characteristic curve (ROC).

#### Enterotype Dysbiosis Score (EDS)

To obtain the enterotype dysbiosis score (EDS), the enterotype strength derived from the FKM clustering for each sample was inverted, z-score normalized, and scaled to have a zero to one scale. As a result, samples with lower enterotype strength in FKM clustering have a higher dysbiosis score. The dysbiosis score by Lloyd-Price et al. 2019 was calculated as described in the original study^6^ using the samples in our cohort with a reported absence of disease as the healthy reference cohort.

The Kaplan-Meier survival analysis was done using the *surv* fuction to calculate the survival time and the *survfit* function to fit the Kaplan-Meier curve from the *survival* R package^67^. P-values were calculated using a log-rank test.

#### Enterotype prediction for novel samples

XGboost regression models^43^ were constructed to predict the strength of each enterotype, which was determined by FKM clustering. The prediction models were trained based on the genus-level taxonomic profiles of the dataset using the *train* function of the caret package in R^68^ with a 10-times repeated 10-fold cross-validation procedure. Before the model construction, minor genera with an average relative abundance of <1e-4 were excluded, and only abundant genera above the threshold were used for the training. To avoid model overfitting due to multiple samples derived from the same individuals, we ensured these samples were incorporated exclusively in the training or the evaluation data during each cross-validation fold. The models’ accuracies were evaluated by applying the prediction models to the unused validation dataset (347 samples from three studies). Separate models were constructed for each enterotype (i.e., two models for the two-enterotype clustering and three models for three-enterotype clustering). The highest strength obtained from these models was used for each sample’s enterotype classification and EDS.

Additionally, binary LASSO classification models were constructed to predict enterotypes determined by PAM clustering. The models were trained and built using the same procedure described above. The highest score obtained from the LASSO classification models was used for enterotype classification.

The Enterotype classifications and EDS prediction for new independent datasets are offered via an online and publicly accessible “Enterotyper” at https://enterotype.embl.de (Figure 5, C). Users will receive the enterotype classifications as tabular data after submitting taxonomic profiles to the “Enterotyper,” which accepts data from various taxonomic profiling tools. Default settings use the three-enterotype FKM model, which will also report the enterotype classification strength and the EDS. The EDS is calculated by inverting and z-score normalizing using the center and standard deviation from the global enterotype training data as a reference and scaling the maximum strength for an enterotype for each sample. However, XGBoost models reflecting different clustering algorithms (FKM, PAM) and numbers of enterotypes (2, 3, 4) are also provided to allow additional explorative analysis by informed users. Furthermore, a first graphical overview of the classified enterotypes and EDS on the users’ input sample will be displayed compared to the global dataset from this study.

## Data Availability

The MICROB-PREDICT cohort dataset is available in the European Nucleotide Archive and can be accessed via BioProject PRJEB89898 (https://www.ebi.ac.uk/ena/browser/view/PRJEB89898) and PRJEB89899 (https://www.ebi.ac.uk/ena/browser/view/PRJEB89899). The Global Fecal Metagenome dataset, consisting of studies reported in Supplementary Table 1, which are reused in this study and have been extracted from the SPIRE resource^53^ (https://spire.embl.de/). The contextual data (curated for this study), the R code for enterotype clustering, statistical analysis, and building the Enterotyper models are available on GitHub (https://github.com/grp-bork/enterotypes). The Enterotyper is a prediction tool designed for the scientific community to classify human fecal metagenomes into established enterotypes. It can be found at https://enterotype.embl.de.

## Acknowledgments

We thank all members of the Bork group, Jeroen Raes and Jorge Francisco Vázquez Castellano from the Raes group, and all members of the MICROB-PREDICT study group, especially Mani Arumugam, for fruitful discussions. We are grateful to Julien Tap and Falk Hildebrand for their help and support of open science. We also thank Anna Głazek, Anna Schwarz, Roman Thielemann, Leonie Thomas, Ela Cetin, and Moritz von Stetten for their help with the metadata curation.

## Ethics declarations

### Ethics approval and consent to participate

The MUCOSA-PREDICT study (ClinicalTrials.gov number, NCT03056612) is a European, investigator-initiated, multicenter, prospective, observational study^56^. The study was approved by the local ethics committee (35281-2/2017/EKU) in all 48 participating universities, and all patients signed an informed written consent following the Helsinki Declaration.

### Consent for publication

Not applicable.

## Author Contributions

Conceptualization, M.I.K. T.V.R, M.K., P.B.; methodology, M.I.K, T.V.R, M.K., S.N., D.P., F.M.; data analysis, M.I.K, T.V.R., C.Y.K., S.N., J. R., T.S.S, A.O.; collection of samples, metagenomes and metadata curation, T.S.S., A.N.F., W.A., M.K., A.T., R.H., D.H.M.H., R.S., S.K., M.J.P., W.G., F.E.U., K.H.T., S.J., M.T.; Enterotyper prediction tool, M.I.K, S.N, S.M.R, D.P, C.Y.K, C.S, A.F, I.L, T.V.R, M.K; writing, M.I.K, T.V.R, M.K., D.P., T.S.S., S.N., D.P., P.B.; supervision, T.V.R, M.K., D.P., J. T., P.B. All authors discussed the results, reviewed the manuscript, and approved the final manuscript.

## List of abbreviations

AUC: Area under the curve
CH-index: Calinski-Harabasz index
DMM: Dirichlet multinomial mixtures
EDS: Enterotype dysbiosis score
FKM: Fuzzy *K*-means clustering
glm: generalized linear model
GTDB: Genome taxonomy database
LASSO: Least absolute shrinkage and selection operator
JSD: Jenson-Shannon divergence
KEGG: Kyoto encyclopedia of genes and genomes
lmer: linear mixed effect model
MELD: model of end-stage liver disease
mOTU: marker gene-based operational taxonomic unit
NCBI: National center for biotechnology information
PAM: partition around medoid clustering
PCA: Principal components analysis
PCoA: Principal coordinate analysis
PERMANOVA: permutational multivariate analysis of variance
PHATE: Potential of heat-diffusion for affinity-based trajectory embedding
TE: Transient Elastography
UMAP: Uniform manifold approximation and projection
XGBoost: Extreme gradient boosting

