## SupplementaryFigures for "Refined Enterotyping Reveals Dysbiosis in Global Fecal Metagenomes"

### Supplementary Figures (Keller et al. Enterotypes)

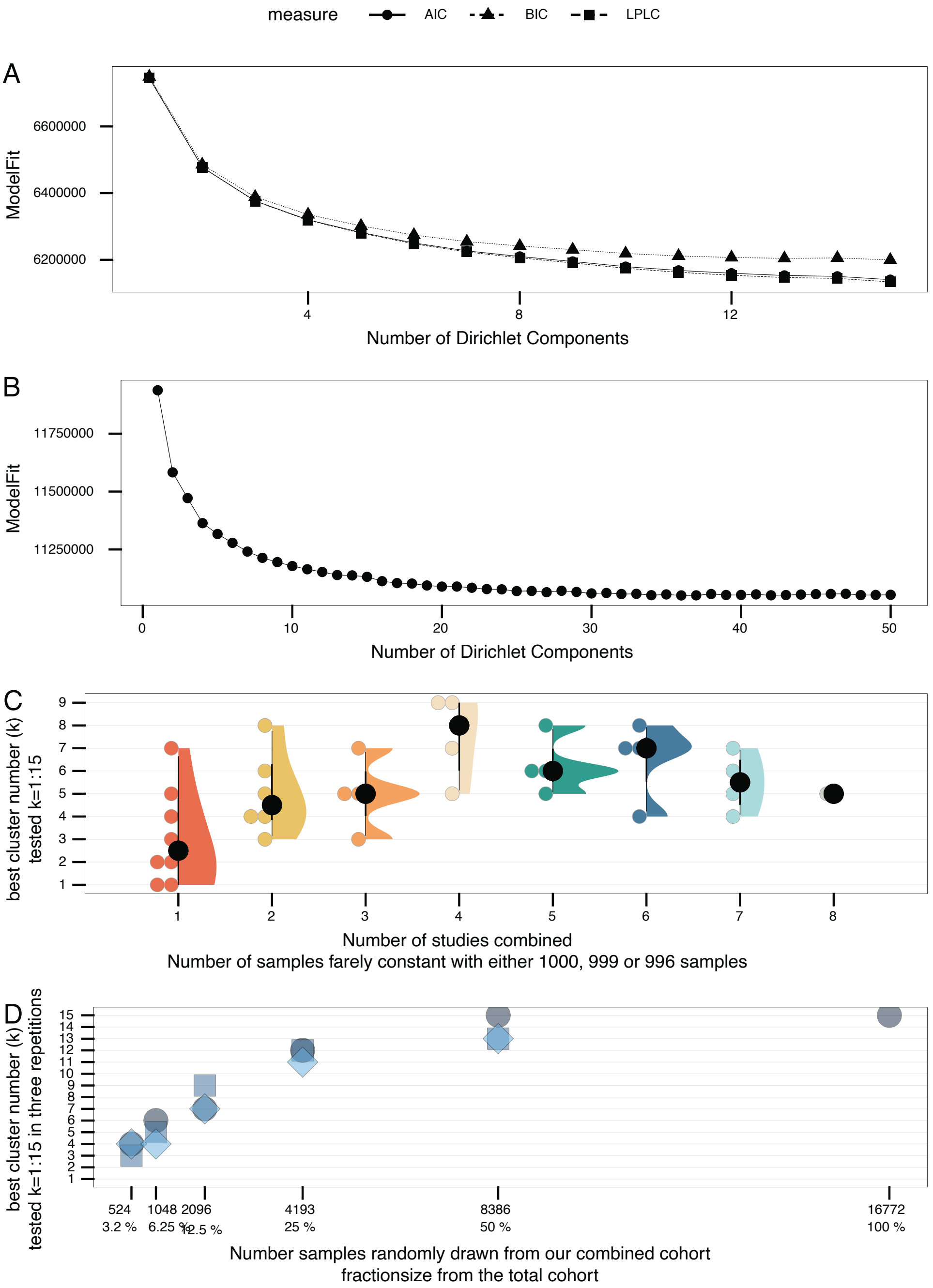


**Figure S1: DMM Clustering. A.** The DMM clustering on the total cohort showed no local minimum using the Laplace approximation, AIC, or BIC when evaluating the clusters up to 15 clusters. **B.** The Laplace approximation was also calculated for DMM cluster up to 50 cluster. **C.** Benchmarking the DMM clustering with increasing heterogeneity by adding samples from increasing numbers of studies in equal proportions. The number of samples was kept almost the same throughout the benchmarking. **D.** Benchmarking the DMM clustering with an increasing number of samples randomly picked from our cohort to keep a similar level of heterogeneity. The selection process was repeated three times.

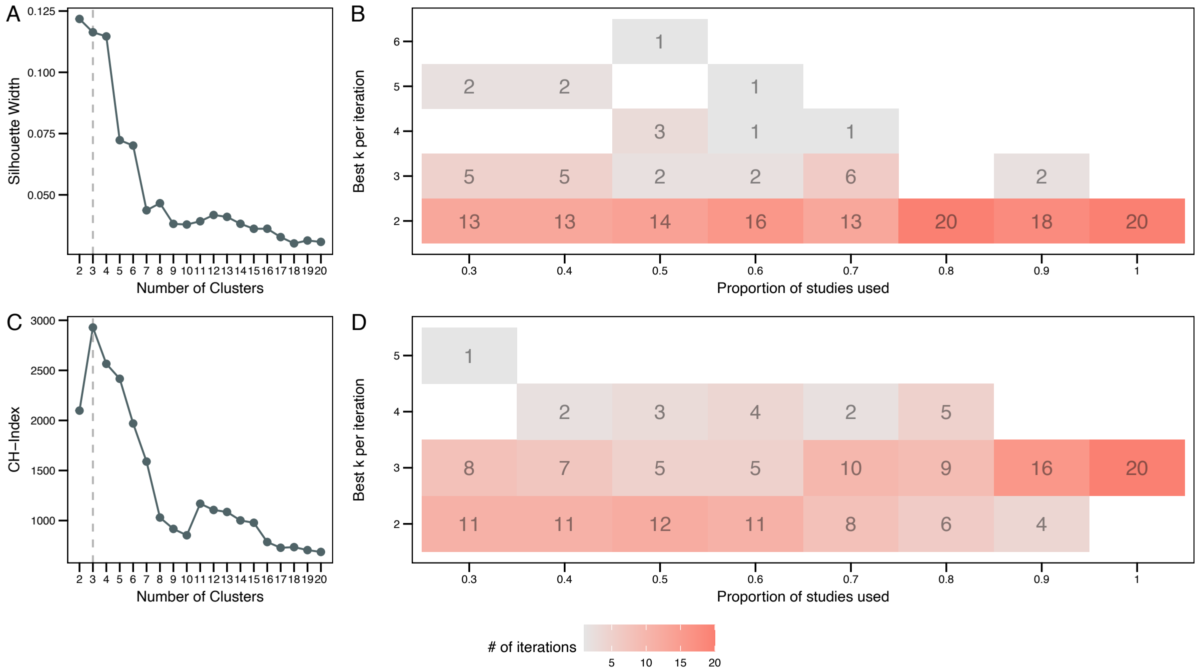


**Figure S2: Statistics to determine the optimal number of enterotypes using the PAM clustering. A + C.** Silhouette width (A) and CH-index (B) for up to 20 possible clusters **B + D.** Rarefaction analysis showing the optimal number of clusters determined by silhouette width (B) and CH-Index (C) for samples from a subset of studies representing 30% - 100% of the studies. This selection process was repeated 20 times for each percentage (except 100%), and the PAM clustering workflow was performed on the subset. The best cluster number, k, was chosen by selecting the cluster number that resulted in the highest CH-index or silhouette width, respectively.

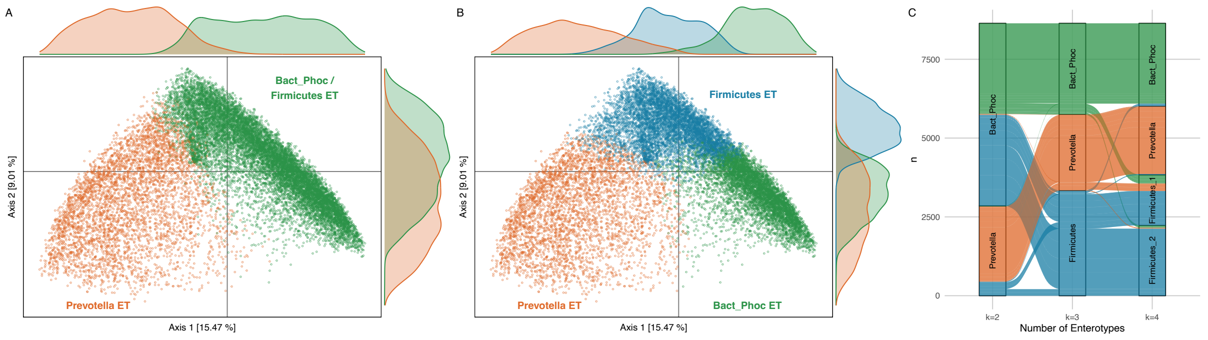


**Figure S3: Visualizing enterotype classifications using PAM clustering. A-B.** Principal coordinates analysis (PCoA) plots derived from the JSD matrix based on genus-level profiles. Samples are colored according to their enterotype classification, as determined by two enterotypes (A) and three enterotypes (B) using PAM clustering. Density plots illustrate the distribution of points along the first two principal components for each enterotype. **C.** Alluvial diagram depicting the consistency of enterotype classification for each sample across the two, three-, and four-enterotype PAM clustering models. Each line represents one sample and is colored based on the three-enterotype model.

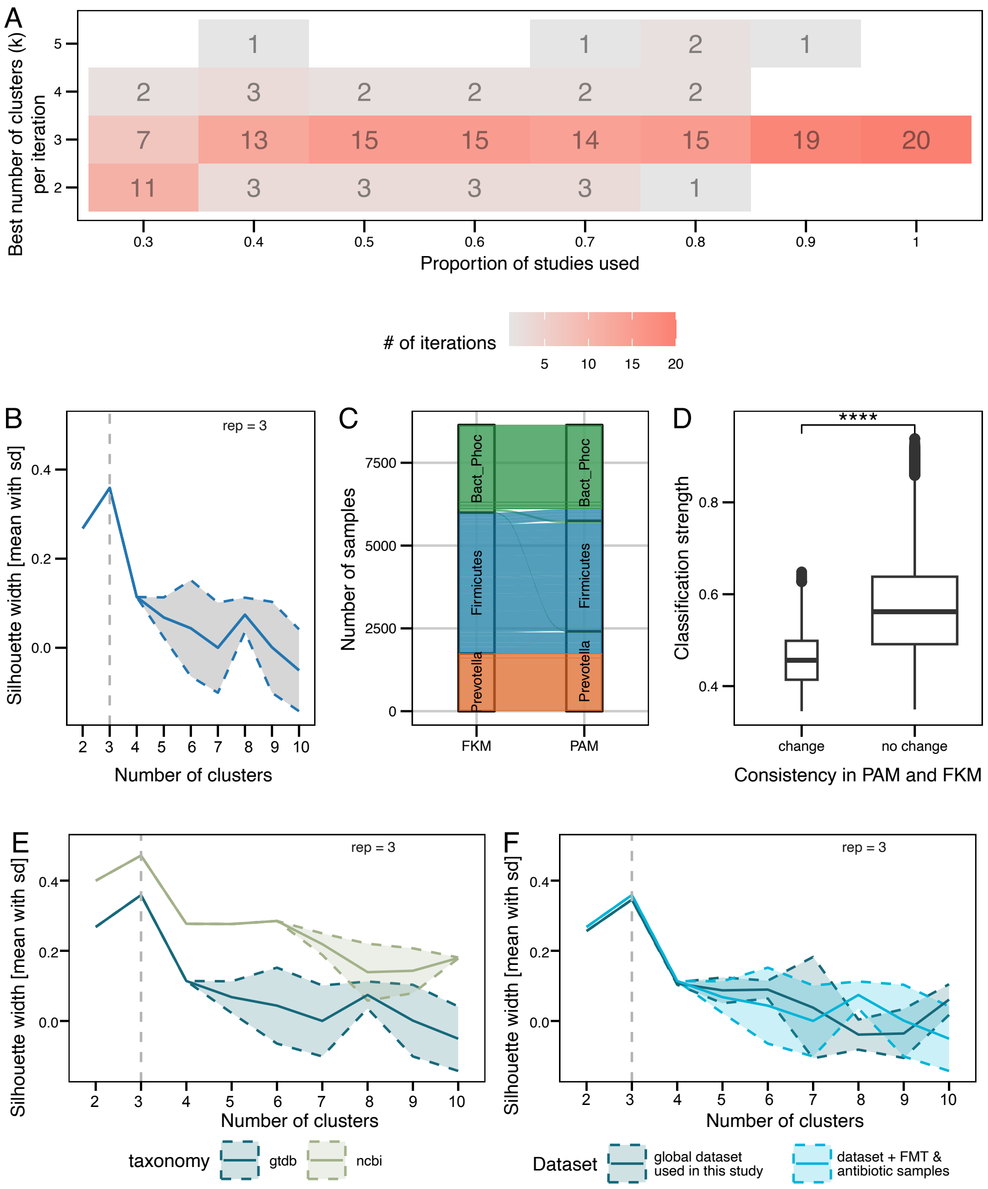


**Figure S4: Statistics to determine the optimal number of enterotypes using the FKM clustering.** **A.** Rarefaction analysis showing the optimal cluster number for samples from subsets of studies representing 30% - 100 % of the studies. This selection process and FKM clustering were repeated 20 times for each percentage (selection in 100% cohort not applicable). The best cluster number was chosen by selecting the number that resulted in the highest silhouette width **B.** Silhouette width mean values with confidence intervals for different cluster numbers tested by the FKM algorithm. **C.** Alluvial diagram demonstrating the overlap between enterotype classifications derived from FKM and PAM clustering methods. Each line represents one sample and is colored according to the three-enterotype FKM clustering model. **D.** Classification strength derived from the FKM clustering comparing samples that change their enterotype classification between FKM and PAM clustering and samples that do not. **E.** Silhouette width derived from the FKM clustering on GTDB and NCBI based taxonomic profiles. **F.** Silhouette width derived from the FKM clustering on the global dataset and on the dataset together with samples with associated antibiotic usage or fecal microbiome transplantation (FKM)

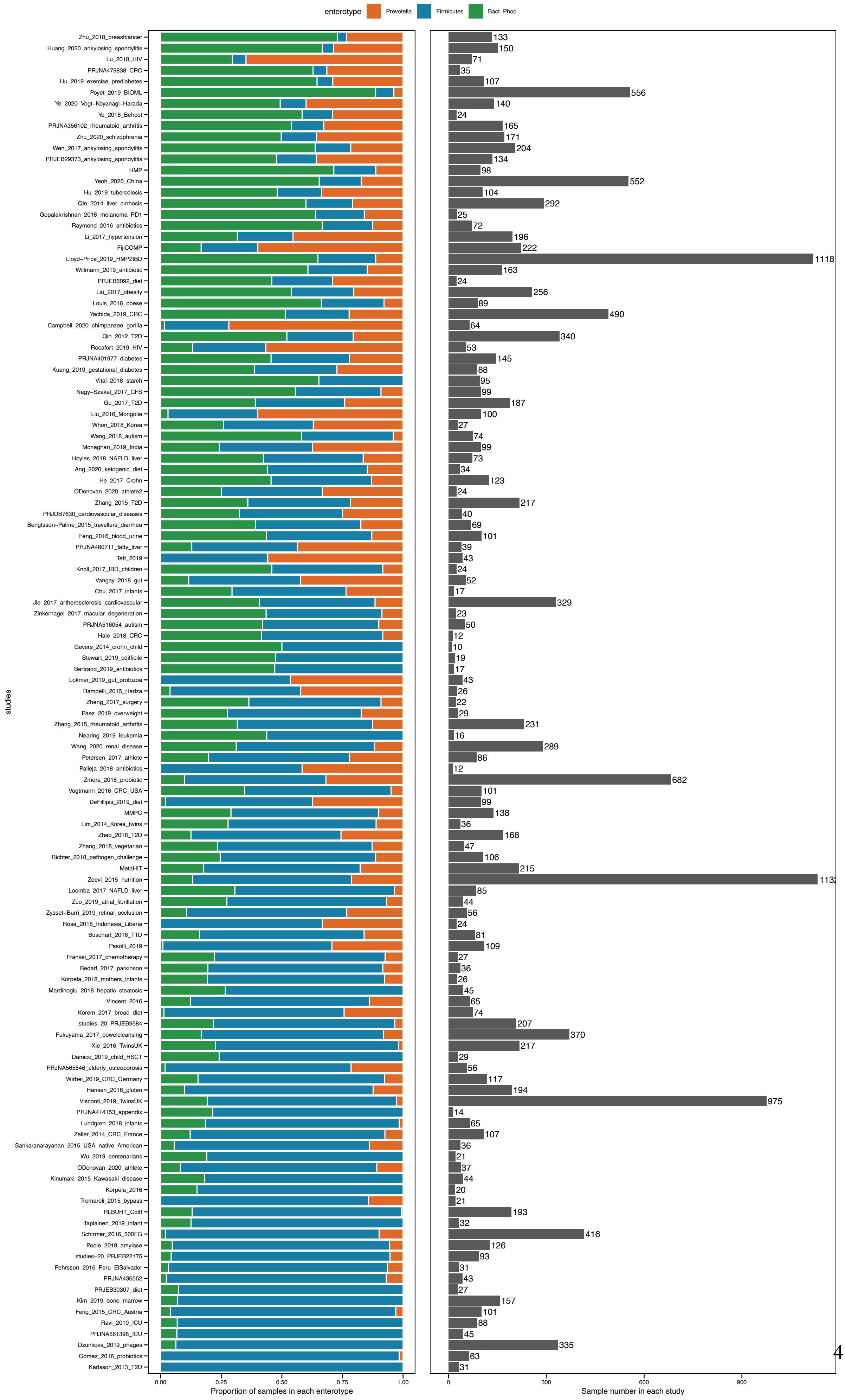
enterotype
Prevotella
Firmicutes
Bact_Phoc
studies
Zhu_2018_breastcancer
Huang_2020_ankylosing_spondylitis
Lu_2018_HIV
PRJNA479838_CRC
Liu_2019_exercise_prediabetes
Poyet_2019_BIOML
Ye_2020_Vogt−Koyanagi−Harada
Ye_2018_Behcet
PRJNA356102_rheumatoid_arthritis
Zhu_2020_schizophrenia
Wen_2017_ankylosing_spondylitis
PRJEB29373_ankylosing_spondylitis
HMP
Yeoh_2020_China
Hu_2019_tubercolosis
Qin_2014_liver_cirrhosis
Gopalakrishnan_2018_melanoma_PD1
Raymond_2016_antibiotics
Li_2017_hypertension
FijiCOMP
Lloyd−Price_2019_HMP2IBD
Willmann_2019_antibiotic
PRJEB6092_diet
Liu_2017_obesity
Louis_2016_obese
Yachida_2019_CRC
Campbell_2020_chimpanzee_gorilla
Qin_2012_T2D
Rocafort_2019_HIV
PRJNA401977_diabetes
Kuang_2019_gestational_diabetes
Vital_2018_starch
Nagy−Szakal_2017_CFS
Gu_2017_T2D
Liu_2016_Mongolia
Whon_2018_Korea
Wang_2018_autism
Monaghan_2019_India
Hoyles_2018_NAFLD_liver
Ang_2020_ketogenic_diet
He_2017_Crohn
ODonovan_2020_athlete2
Zhang_2015_T2D
PRJDB7630_cardiovascular_diseases
Bengtsson−Palme_2015_travellers_diarrhea
Feng_2016_blood_urine
PRJNA480711_fatty_liver
Tett_2019
Knoll_2017_IBD_children
Vangay_2018_gut
Chu_2017_infants
Jie_2017_artherosclerosis_cardiovascular
Zinkernagel_2017_macular_degeneration
PRJNA516054_autism
Hale_2018_CRC
Gevers_2014_crohn_child
Stewart_2019_cdifficile
Bertrand_2019_antibiotics
Lokmer_2019_gut_protozoa
Rampelli_2015_Hadza
Zheng_2017_surgery
Paez_2019_overweight
Zhang_2015_rheumatoid_arthritis
Nearing_2019_leukemia
Wang_2020_renal_disease
Petersen_2017_athlete
Palleja_2018_antibiotics
Zmora_2018_probiotic
Vogtmann_2016_CRC_USA
DeFillipis_2019_diet
MMPC
Lim_2014_Korea_twins
Zhao_2018_T2D
Zhang_2018_vegetarian
Richter_2018_pathogen_challenge
MetaHIT
Zeevi_2015_nutrition
Loomba_2017_NAFLD_liver
Zuo_2019_atrial_fibrillation
Zysset−Burri_2019_retinal_occlusion
Rosa_2018_Indonesia_Liberia
Buschart_2016_T1D
Pasolli_2019
Frankel_2017_chemotherapy
Bedarf_2017_parkinson
Korpela_2018_mothers_infants
Mardinoglu_2018_hepatic_steatosis
Vincent_2016
Korem_2017_bread_diet
studies−20_PRJEB9584
Fukuyama_2017_bowelcleansing
Xie_2016_TwinsUK
Damico_2019_child_HSCT
PRJNA565546_elderly_osteoporosis
Wirbel_2019_CRC_Germany
Hansen_2018_gluten
Visconti_2019_TwinsUK
PRJNA414153_appendix
Lundgren_2018_infants
Zeller_2014_CRC_France
Sankaranarayanan_2015_USA_native_American
Wu_2019_centenarians
ODonovan_2020_athlete
Kinumaki_2015_Kawasaki_disease
Korpela_2016
Tremaroli_2015_bypass
RLBUHT_Cdiff
Tapiainen_2019_infant
Schirmer_2016_500FG
Poole_2019_amylase
studies−20_PRJEB22175
Pehrsson_2016_Peru_ElSalvador
PRJNA436562
PRJEB30307_diet
Kim_2019_bone_marrow
Feng_2015_CRC_Austria
Ravi_2019_ICU
PRJNA561398_ICU
Dzunkova_2019_phages
Gomez_2016_probiotics
Karlsson_2013_T2D
0.00
0.25
0.50
0.75
1.00
Proportion of samples in each enterotype
133
150
71
35
107
556
140
24
165
171
204
134
98
552
104
292
25
72
196
222
1118
163
24
256
89
490
64
340
53
145
88
95
99
187
100
27
74
99
73
34
123
24
217
40
69
101
39
43
24
52
17
329
23
50
12
10
19
17
43
26
22
29
231
16
289
86
12
682
101
99
138
36
168
47
106
215
113
85
44
56
24
81
109
27
36
26
45
65
74
207
370
217
29
56
117
194
975
14
65
107
36
21
37
44
20
21
193
32
416
126
93
31
43
27
157
101
88
45
335
63
31
0
300
600
900
Sample number in each study

**Figure S5**: **The proportion of samples from each study classified to enterotypes.** The stacked bar plots show the distribution of samples classified according to the respective enterotype. The right grey bar indicates the number of samples in that study.

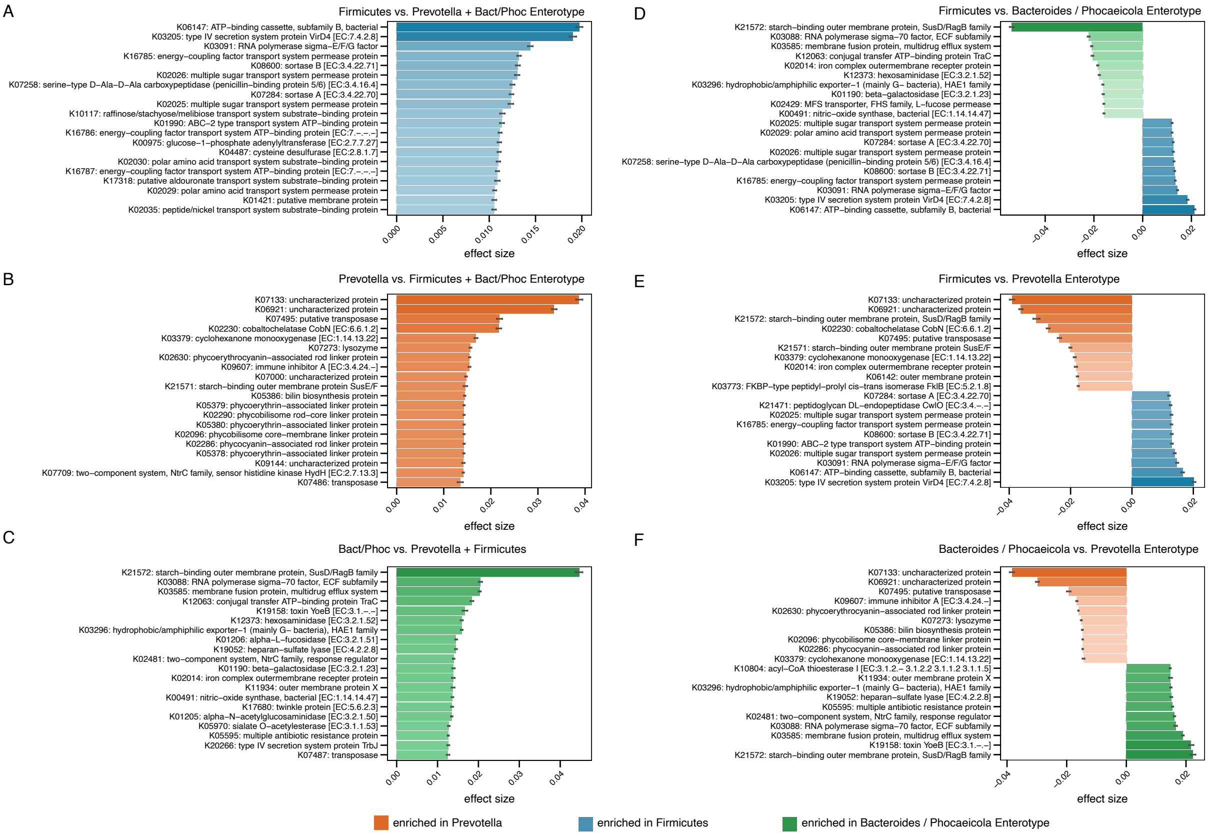


**Figure S6: Functional profiles shown as enriched KEGG-orthologs (KO) for each enterotype.** Effect sizes with confidence intervals derived from differential abundance analysis using the Wilcoxon test with Benjamini-Hochberg p-value adjustment for false discovery rate on the relative abundances of KOs. We compared the samples classified with each enterotype with those not classified into this enterotype (**A-C**). In addition, and inspired by the sample distribution in the MDS plot (Figure 2D), we compared the KO abundances between the Firmicutes enterotype with the Bacteroides/Phocaeicola (**D**) or Prevotella (**E**) enterotype and directly the Bacteroides/Phocaeicola with the Prevotella enterotype (**F**). The colors of the bars represent the enterotype in which this KO shows an enrichment. Only the top KO with significant changes reported by an adjusted p-value < 0.01 are shown.

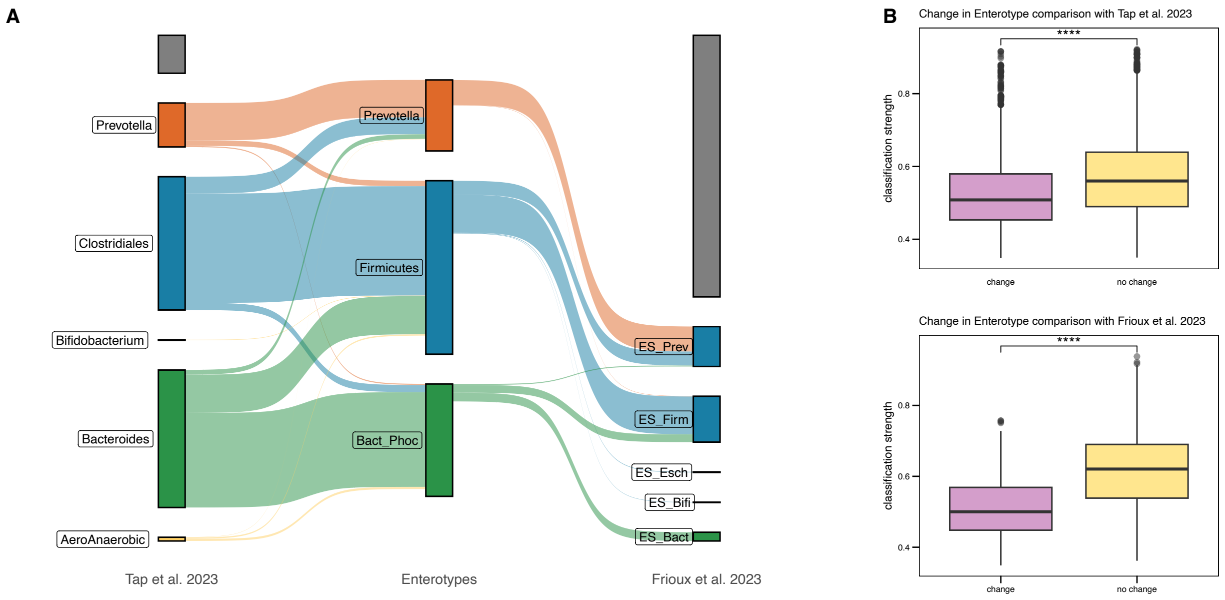


**Figure S7: Enterotypes comparison with Tap et al. 2023 and Frioux et al. 2023. A.** Comparison of the Enterotype classification from this study with the overlapping samples from Tap et al. 2023 (left, n = 5,668) and Frioux et al. 2023 (right, n = 1,812). The grey bars on the top left indicate the number of samples included in this study, but not be Tap or Frioux, respectively. **B.** Samples that changed their enterotype assignment showed lower FKM-derived classification strength. **** $p \leq 0.0001$.

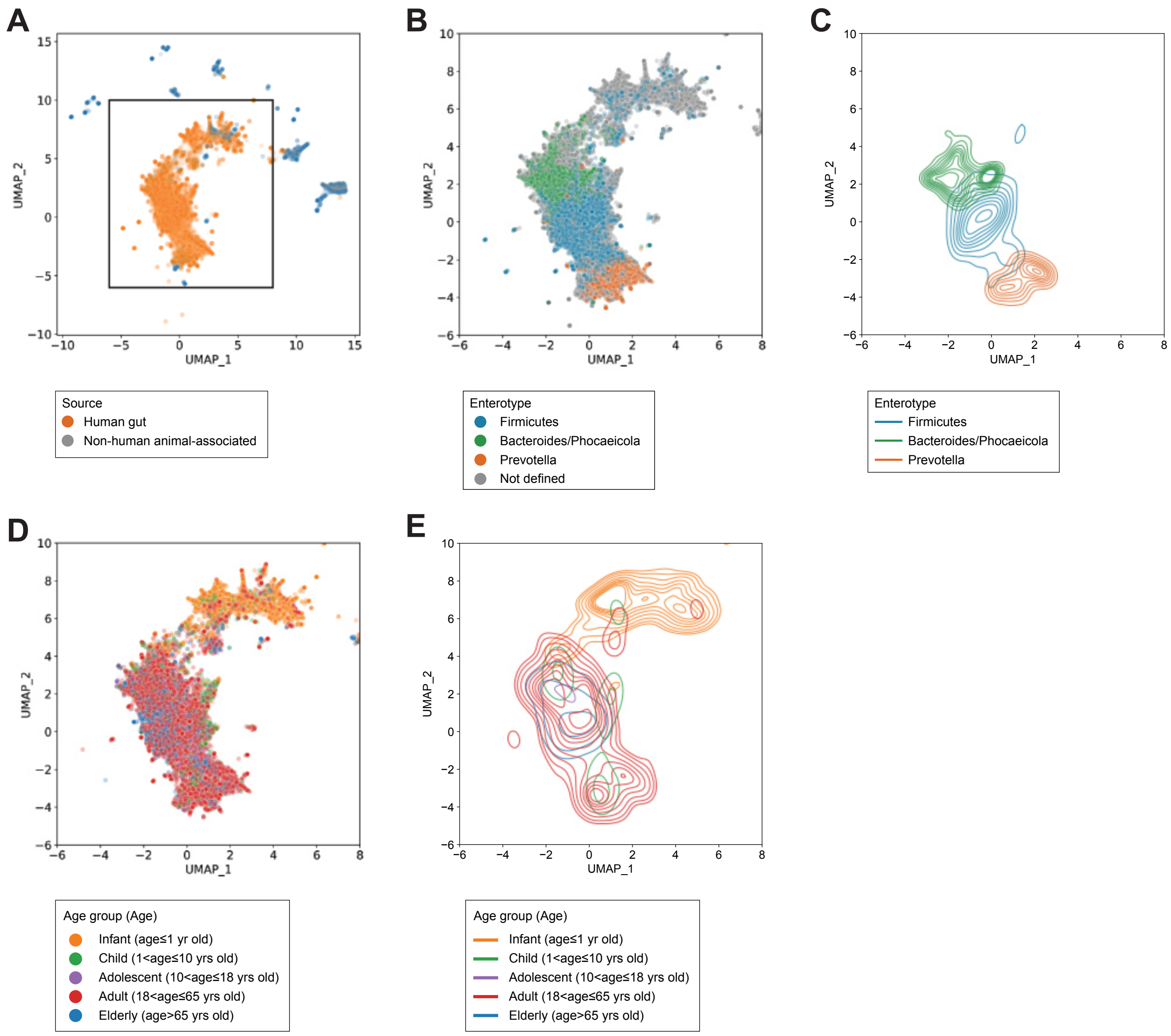


**Figure S8: Imposing of enterotypes on UMAP based on the global dataset together with other microbiome sample types A.** UMAP plot of 49,762 metagenomic samples. The black square highlighted the UMAP space where the human fecal samples are enriched. **B**. UMAP plot colored by enterotype of the samples. **C**. Contour plot representing the sample density of each enterotype. **D**. UMAP plot colored by the age group of the human fecal samples. **E**. Contour plot representing the sample density of each age group.

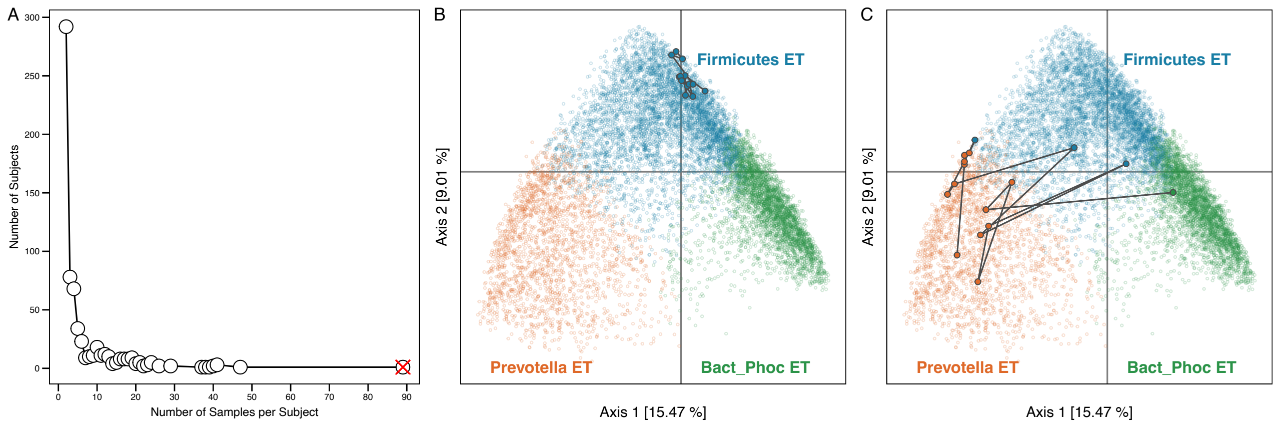


**Figure S9: Analysis of enterotype classification stability A.** The number of subjects with multiple samples. We excluded one subject with 89 samples as an outlier from the analysis using the Markov chain model (Figure 4). **B & C.** Examples of two opposite cases for longitudinal behavior of the subjects' microbiome composition: a very stable example (B) and an unstable (C) case.

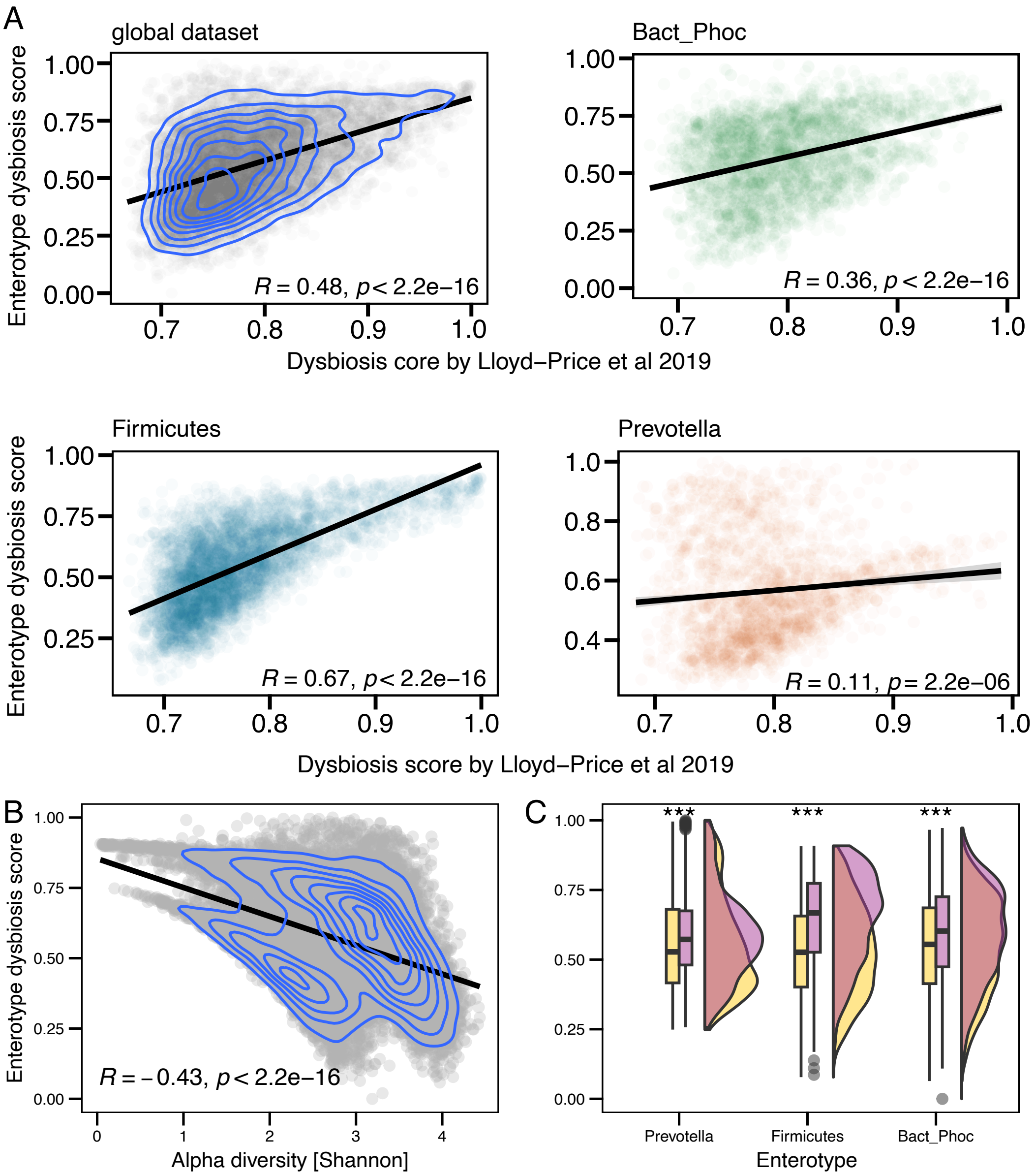


**Figure S10: Enterotype dysbiosis score (EDS) correlates with other dysbiosis estimates. A.** EDS correlates with an already established one dysbiosis score[6] in sample subsets of all three enterotypes. **B.** EDS correlated negatively with alpha diversity **C.** The samples from subjects with a reported disease (n = 3,399) showed a higher enterotype dysbiosis score than those from subjects with no reported disease (n = 8,672) in samples subsets of all three enterotypes. *** $p \leq 0.001$

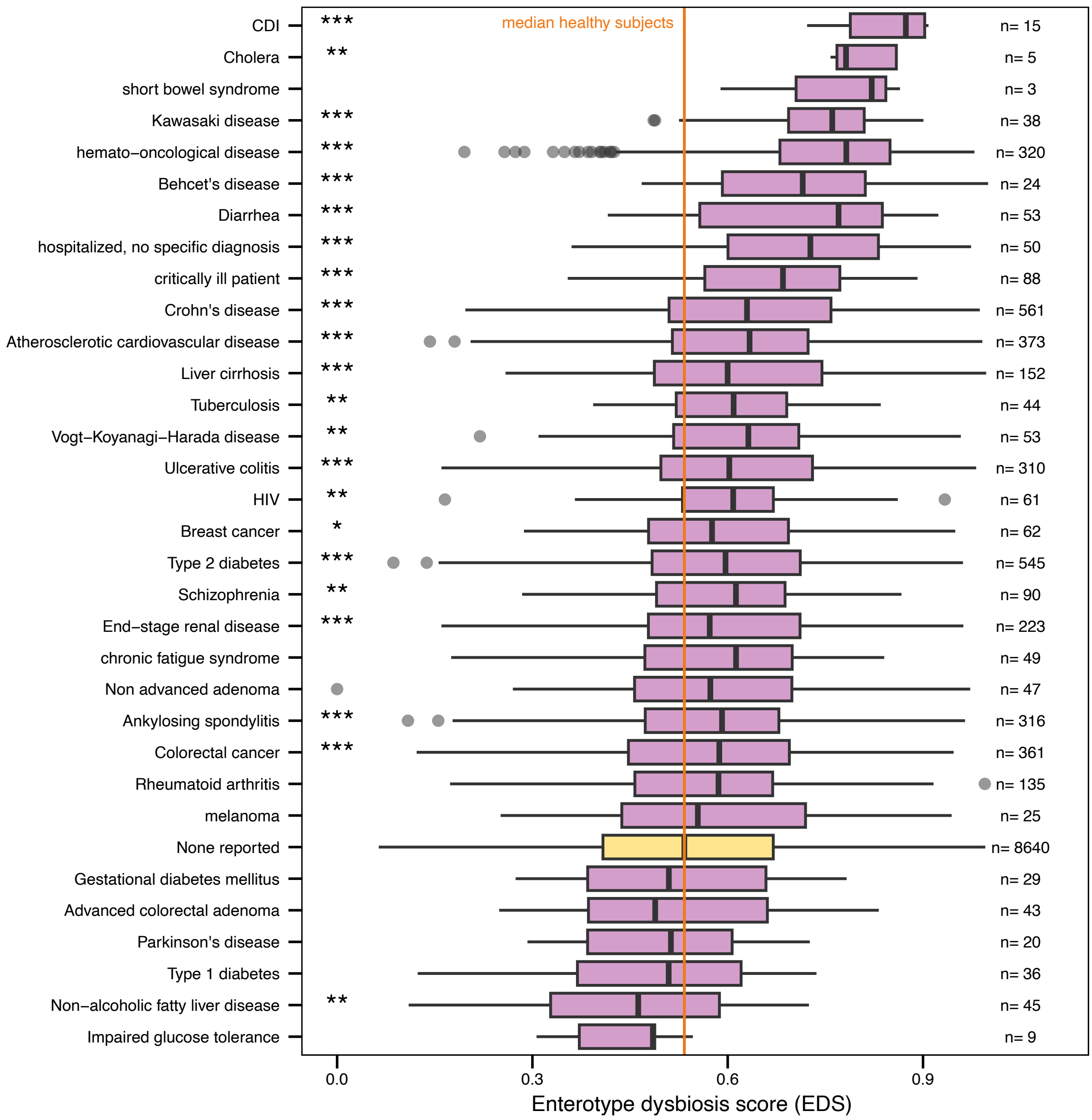


**Figure S11: Enterotype dysbiosis score (EDS) by diseases and symptoms.** The EDS for all samples associated with a disease or symptom compared to the EDS for samples with reported absence of disease (yellow). The difference in EDS from a disease group to the group without disease was calculated using the Wilcoxon test and adjusted for multiple testing using Benjamini-Hochberg correction. $^*q \leq 0.05$, $^{**}q \leq 0.01$, $^{***}q \leq 0.001$.

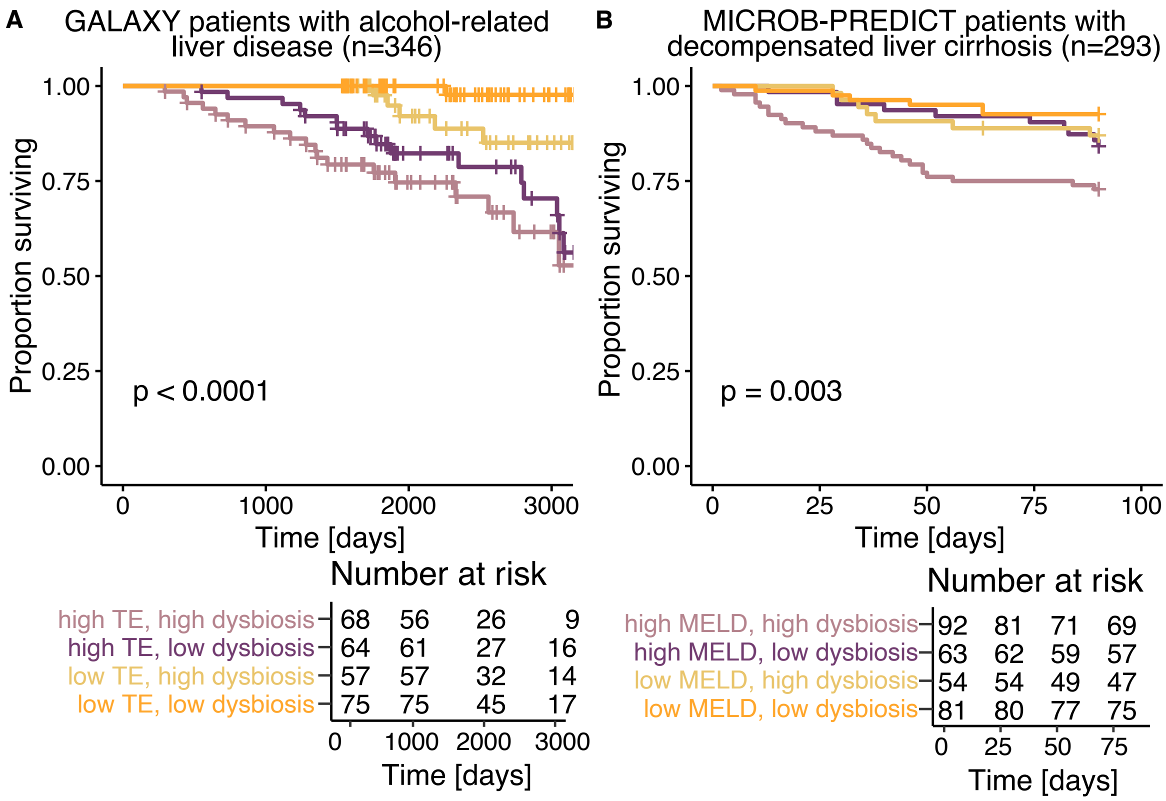


**Figure S12: Enterotype dysbiosis score (EDS) can stratify patients for their disease progression or survival probability. A.** In a cohort of alcohol-related liver disease (ALD) patients, baseline EDS stratifies survivors and non-survivors. TE is a marker to estimate the stiffness of the liver, and subsequently the level of fibrosis. **B.** In patients with decompensated cirrhosis, EDS stratified survivors and non-survivors, especially among those who already have a high MELD (model for end-stage liver disease) score and severe disease. P-values were calculated using a log-rank test.

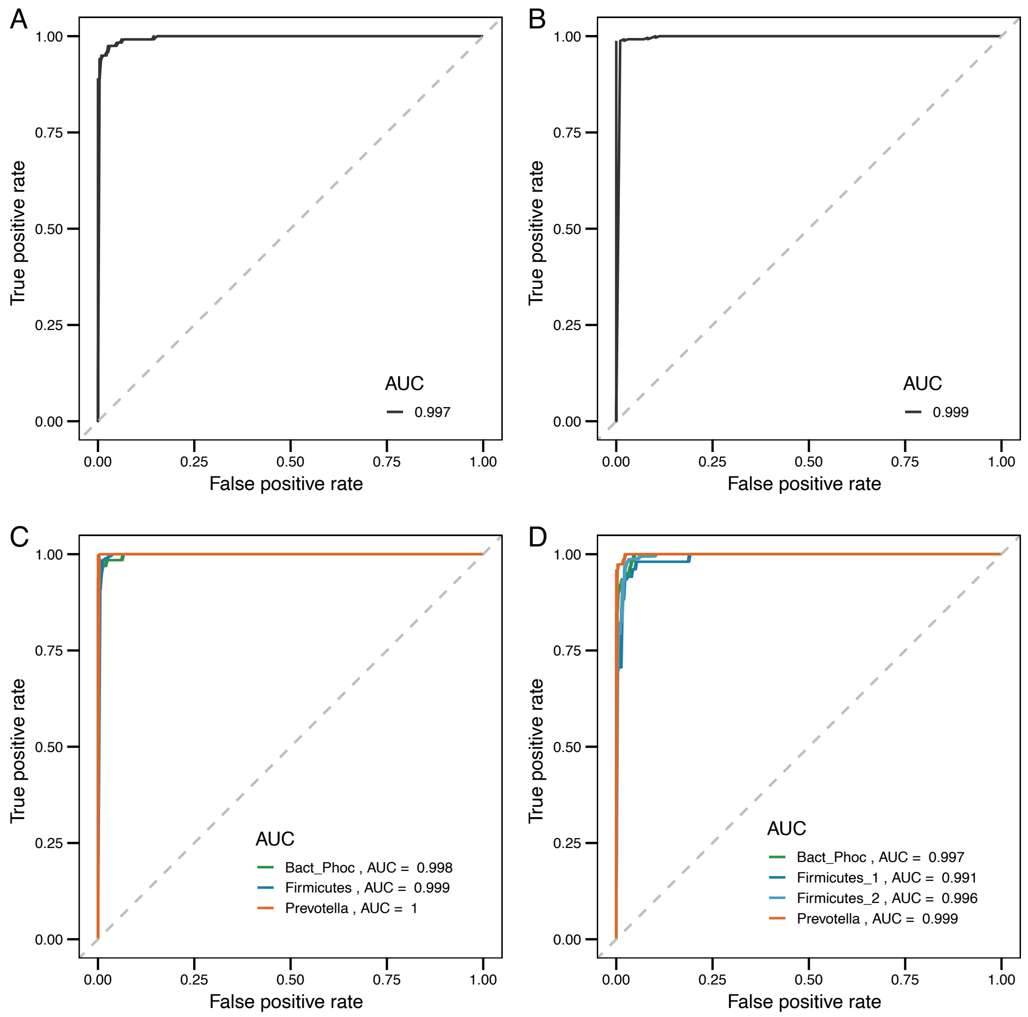


**Figure S13: AUCs for the XGBoost models in the "Enterotyper." A.** The AUC for the FKM-based enterotype classification prediction for the two enterotypes. **B – D**. AUCs for PAM-based enterotype classification prediction for two (B), three (C), and 4 (D) enterotypes. All AUCs have been calculated based on the prediction of the validation dataset.
